# Cryo-EM Structure of the relaxosome, a complex essential for bacterial mating and the spread of antibiotic resistance genes

**DOI:** 10.1101/2024.12.19.629322

**Authors:** Sunanda M. Williams, Sandra Raffl, Sabine Kienesberger, Aravindan Ilangovan, Ellen L. Zechner, Gabriel Waksman

## Abstract

Bacterial mating, or conjugation was discovered nearly 80 years ago as a process transferring genes from one bacterial cell (the donor) to another (the recipient). It requires three key multiprotein complexes in the donor cell: a DNA-processing machinery called the relaxosome, a double-membrane spanning type 4 secretion system (T4SS), and an extracellular appendage termed pilus. While the near-atomic resolution structures of the T4SS and pilus are known, that of the relaxosome is not^1–3^. Here, we describe the cryo-EM structure of the fully assembled relaxosome encoded by the paradigm F plasmid in two different states corresponding to distinct functional steps along the DNA processing reaction. By varying the structures of model DNAs we delineate conformational changes required to initiate conjugation. Mutational studies of the various protein-protein and protein-DNA interaction hubs suggest a complex highly sensitive to trigger signals, that could arise from cell-to-cell contacts with recipient cells.

---

The nucleoprotein complex termed relaxosome contains the relaxase and other auxiliary transfer proteins, which assembles at the “origin of transfer” (*oriT*) region of a plasmid and prepares it for transfer to another bacterial cell^4,5^. This process whereby plasmids spread among bacterial populations through cell-to-cell contact is termed bacterial conjugation, an important way to exchange genetic material and plays a crucial role in bacterial adaptation and evolution^6^. It is also one of the most effective means by which antibiotic resistance genes spread among bacterial populations resulting in antimicrobial resistance (AMR)^7–11^. AMR is considered one of the most severe global public health and development threats of this century^12^. F plasmid which we are focusing here, are the main carriers of antibiotic resistance genes in *E. coli* and historically associated with resistance or ‘R factors’^13^.

All DNAs transferred via conjugation do so from an essential sequence called *oriT*, a stretch of DNA about 400 base pairs (bp) in the F plasmid^14–16^. The F plasmid relaxosome complex contains, in addition to *oriT*, three plasmid-encoded proteins including the DNA-processing enzyme TraI (generally known as the “relaxase”) and 2 accessory proteins TraY and TraM, and IHF, a protein encoded by the bacterial genome (ED Fig. 1a). To date, multiple binding sites for these proteins have been characterised on *oriT* (2 TraI binding sites (*traI_helicase_* and *traI_TE_*), 2 IHF binding sites (*IHFa,b*), 3 TraM binding sites (*sbmA,B,C*) and 2 TraY binding sites (*sbyA,C*); details in ED Fig. 1b).

**Fig. 1.**
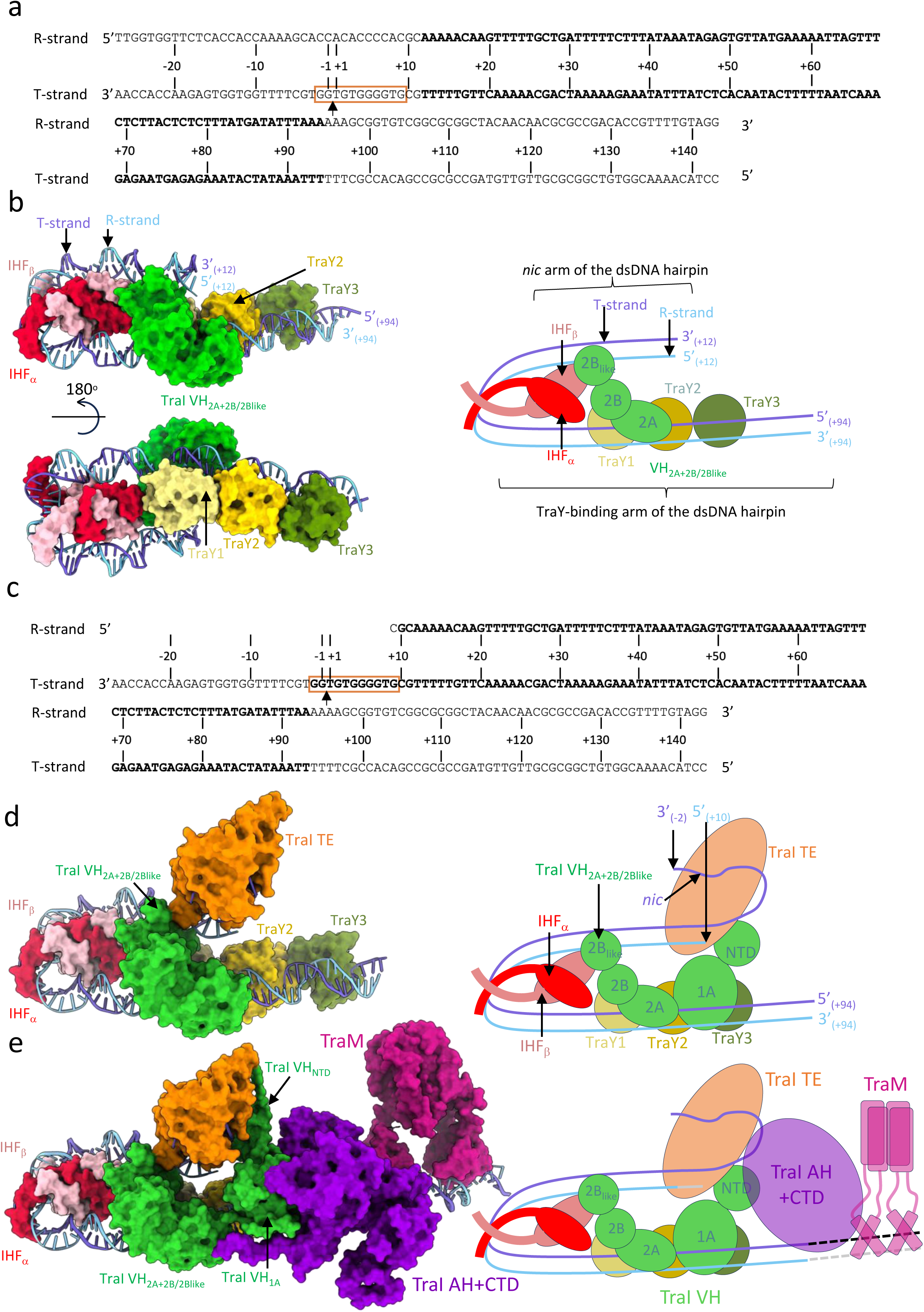
Structures of the ds_-27_+143_-R relaxosome, the ss_-27_+8_ds_+9_+143_-R relaxosome, and the fully-assembled ss_-27_+8_ds_+9_+143_-R relaxosome. a, DNA used to solve the structure of ds_-27_+143_-R. Nomenclature and numbering is as in ED Fig. 2c. *nic* is indicated by an arrow. The region of DNA for which a model could be derived from the density is shown in bold. Replicative (R-strand) and transfer (T-strand) strands are indicated. Orange box locates the TE-binding site on the T-strand between bp_-2_ and bp_+9_. b, Structure of ds_-27_+143_-R. This structure was derived from the “locally-refined” map shown in ED Fig. 5a. Left: Two orientations of the complex. Proteins and DNA are shown in surface and ribbon, respectively, colour-coded dark and light blue for the T- and the R-strand respectively; red and pink for IHFα and IHFβ, respectively; pale yellow, bright yellow and olive green for TraY1, TraY2, and TraY3, respectively; and green for the only TraI part visible in the density for this complex, VH_2A+2B/2B-like_. 5’ and 3’ end base pairs are labelled. Right: Schematic diagram of the structure in the orientation shown at upper left. The two arms of the dsDNA hairpin are labelled. Same colour-coding as at left is used. c, DNA used to solve the structures of ss_-27_+8_ds_+9_+143_-R complex (ED Fig. 2c). *nic*, the region of DNA for which a model could be derived from the density, and the location of the TE-binding site on the T-strand are shown as in panel a. d, Structure of ss_-27_+8_ds_+9_+143_-R. This model is derived from the locally-refined map described in ED Fig. 5b. Proteins and DNA are shown in surface and ribbon, respectively, colour-coded as in panel b, except for the additional TE domain in orange. Left: Near-atomic resolution structure of the complex. Right: Schematic representation of the structure. 3’ and 5’ DNA ends are indicated as well as the location of the *nic* site. e, Structure of the fully-assembled ss_-27_+8_ds_+9_+143_-R relaxosome. This model is derived from both the locally-refined and global maps reported in ED Figs. 5b-d and 5h-j. Proteins and DNA are shown in surface and ribbon, respectively, colour-coded as in panel d. TraI AH+CTD is in violet and TraM is in magenta. Left: model of the complex. Right: Schematic diagram.

TraI is a multi-domain protein that consists of an N-terminal trans-esterase (TE) domain that contains its “relaxase” activity, followed by two helicase domains (one vestigial (VH), the other active (AH)), and a C-terminal domain (CTD) responsible for interactions with TraM (ED Fig. 1c,d)^17–21^. IHF, TraY, and TraM are non-catalytic proteins. IHF, a binary complex of two very similar proteins, IHFα and IHFβ, is known to induce a sharp U-turn bend on its binding site (ED Fig. 1e)^22^. TraM is responsible for binding to both its cognate *oriT* regions and to the C-terminal tail of TraD (ED Fig. 1f)^23,24^, a T4SS protein that functions as recruitment platform for the relaxosome. TraY has DNA-bending ability and is known to stimulate TraI activity when bound to *oriT* ^25^. Once formed, the relaxosome is recruited to the T4SS and the relaxosome-T4SS super-complex may rest quiescent until contacts between donor and recipient cells are established, activating the relaxosome. Activation initiates a yet uncharacterised cascade of molecular events resulting in the nicking of one of the plasmid DNA strands (the T-strand to be transferred) at a site called “*nic*” (ED Fig. 1b) by the TE activity of TraI. The TraI catalysed nicking is stimulated by the other proteins in the relaxosome^25,26^. Nicking is followed by covalent attachment of TraI to the free 5’ phosphate resulting from the nicking reaction yielding a protein-ssDNA conjugate that, once unwound from its complementary strand (the R strand), is transported through the T4SS into the recipient cell^3,27–29^. To achieve this, another molecule of TraI functions as a helicase, unwinding the dsDNA in the 5’ to 3’ direction^17,30,31^. Once transfer is completed, both the transferred T-strand and the copy retained by the donor (the R-strand) undergo replication to duplex DNA thereby increasing the abundance of the transferred genome (and any antibiotic resistance genes it may carry) within a bacterial population^32^.

A fully functional assembly of the relaxosome is crucial for effective nicking and transfer of the plasmid. Mutation of a catalytic tyrosine in the TraI active site reduces plasmid transfer by ∼ 5000-fold. *oriT* was shown to have functional domains which individually contribute to nicking and transfer functions^33^. Deletions of *oriT* binding sites and/or in the absence of other relaxosome proteins, TraI can only bind to ssDNA and cannot initiate transfer from ds *oriT*^31^.

Molecular events controlling the start of conjugative gene transfer are yet to be understood. Some structural information is available on individual protein components of the relaxosome in gram-negative (ED Fig. 1d-f)^17,22,34^ and more recently in gram-positive bacteria^35–38^. However, crucially, there is to date no information on the structure of a fully or even partially assembled relaxosome.

## Assembly of the relaxosome and identification of DNase I footprints of protein components

We first confirmed the feasibility of purifying a fully-assembled relaxosome from purified components. The four proteins were mixed in molar excess of a purified DNA of 316 bp (*oriT*_316_) encompassing all known binding sites for each protein and the resulting complex (*oriT*_316_-R) was purified (ED Fig. 2a,b).

**Fig. 2.**
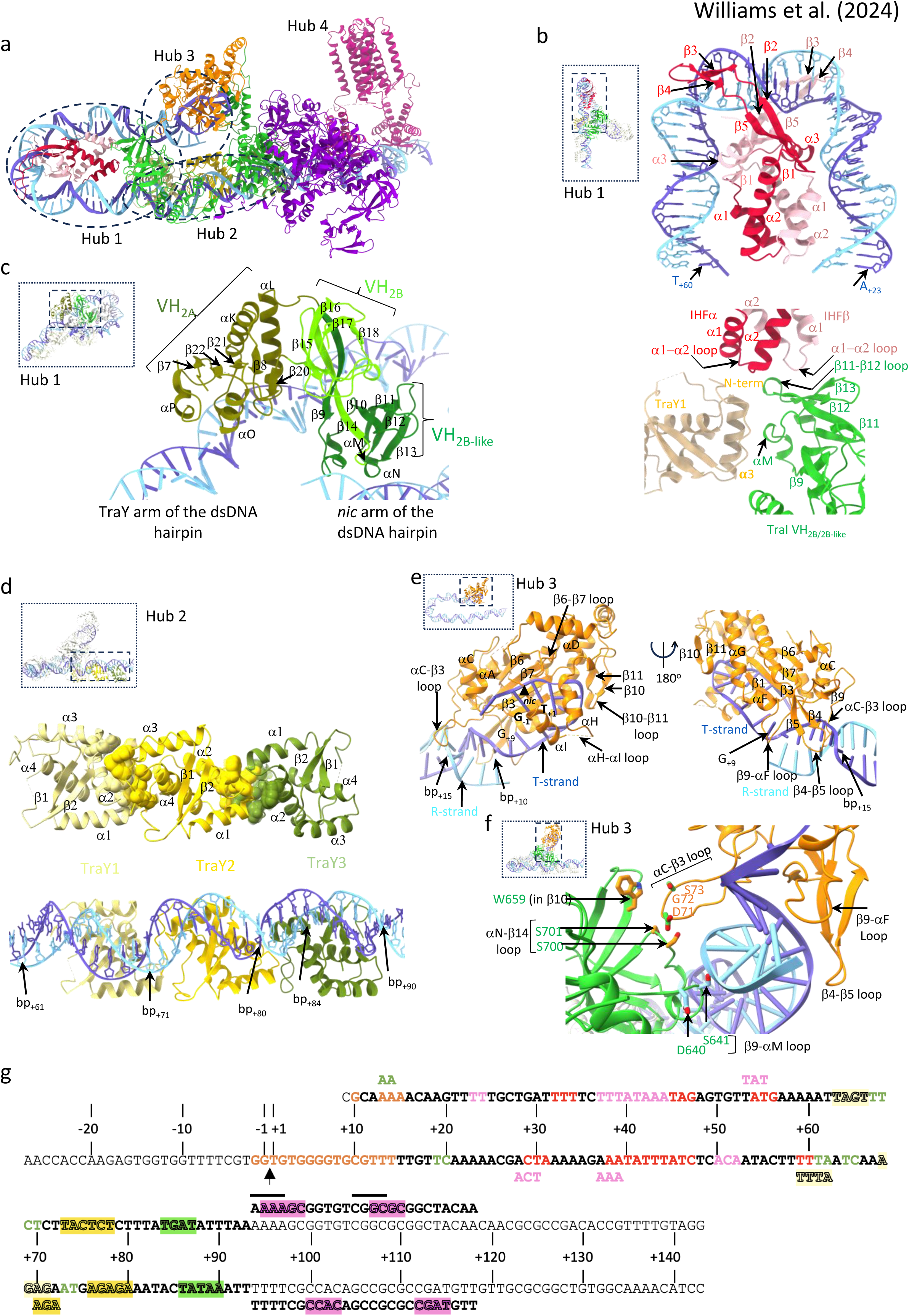
Hub architecture of the ssDNA_-27_+8_ds_+9_+143_-R fully assembled relaxosome structure. Proteins and DNAs are shown in ribbon representation. Secondary structures involved in DNA-binding are labelled. In panels b-f, insets of the entire structure are provided for orientation. Within insets, dash-lined box shows the region(s) detailed in the corresponding panel(s). a, Hub definition. The 4 hubs are shown within dotted lined ovals (except for hub 4) and are labelled hub1-4. b, Hub1. Top: IHF-DNA interaction. Interactions are dominated by residues in the β2-5 region with bending triggered by insertion of the β2-3 hairpin into the major groove of the DNA (see ED Fig. 9a, top panel, for details). Bottom: protein-protein interactions between TraY1, IHF and TraI VH_2A+2B/2B-like_ (see ED Fig. 9a, bottom panel for details). TraY1 has a 12 residue N-terminal tail, the very N-terminus of which is involved in interaction with residues in α1 and the α1-α2 loop of IHFα. The remaining N-terminal tail residues as well as many residues in its α3 helix interact with TraI VH. In TraI VH, residues in αM and the β9-αM and β11-β12 loops are implicated in TraY1 binding while other residues in β12, β13 and in the β11-β12 loop interact with residues in α2 and the α1-α2 loop of both IHFα and β. (see ED Fig. 9a for details) c, Interactions between TraI VH_2A+2B/2B-like_ and DNA. Two regions of TraI VH are responsible for dsDNA binding at *oriT*: the tip of the 2B-like domain and a region of the 2A domain (details in ED Fig. 9b). VH_2B/2B-like_ also forms a three-way interaction in all complexes described here with IHF and TraY1 in hub 1, while also interacting with the TE domain of TraI in the ss_-27_+8_ds_+9_+143_-R complex in hub 3. d, Hub2. Top: protein-protein interactions between TraY molecules. Residues at protein-protein interfaces are shown in spheres. Bottom: TraYs-DNA interactions. Boundary base pairs for each binding site are reported. TraY’s binding to DNA is typical of RHH dimers with the β1 and β2 strands interacting with the major groove of the DNA binding site^47^. But binding of TraY1 and TraY2 (but not TraY3 which is inverted) also extends to the minor groove. TraY1 interaction with TraY2 involves two clusters, but only one is involved in interactions between TraY2 and TraY3 (detailed in ED Fig. 9c). e, Hub3. TraI TE interactions with DNA. Left: view highlighting ssDNA-binding with primarily the β6-β7, αH-αI and β10-β11 loops but also dsDNA-binding, primarily with the αC-β3 loop. Right: view 180° away, highlighting dsDNA interaction with primarily with the αC-β3, β4-β5, and β9-αF loops. The *nic* site is shown as are the binding site boundaries on the DNA (ssDNA between -2 and +9 and dsDNA between +10 and +15). The unpaired +9 base is labelled. f, Hub3 (continued): protein-protein interactions between TraI TE and VH_2A+2B/2B-like_ domains. TraI TE is shown in orange ribbon except the residues interacting with VH_2A+2B/2B-like_ shown in green. VH_2A+2B/2B-like_ is in ribbon and residues interacting with the TE and R-strand are shown in orange and light blue sticks. Residues and secondary structures involved in binding are labelled. One particular loop (αC-β3) stabilises 122 Å^2^ of surface area via interactions with TraI VH_2B/2B-like_ residues in β10 and the αN-β14 loop. g, Footprint of relaxosome proteins on DNA. ssDNA and dsDNA bases involved in binding relaxosome components (including TraM, details in ED Fig. 10c) are colour coded according to the protein they contact (red and pink letters for IHFαβ, green letters for TraI_VH2A+2B/2B-like_, orange letters for TraI TE, and pale yellow, bright yellow, and olive green highlights for TraY1, TraY2, and TraY3, respectively, and magenta highlights for TraM). When two proteins have overlapping binding sites, the sequence is repeated above and coloured accordingly. The TraM footprint is shown above and under the R- and T-strand, respectively, in the *sbmC* region. Solid lines above the R-strand TraM site reports on previous findings based on hydroxyl radical protection assays.

To map the footprint of the fully assembled relaxosome complex, the purified *oriT*_316_-R complex was subjected to DNase I cleavage to hydrolyse regions of *oriT* that are not protected upon complex formation. We show here (ED Fig. 2b) that most protected footprint sequences map to regions of *oriT*_316_ extending from the *nic* site to include *traI_TE_*, *IHFa*, *sbyC*, *sbyA* and partial *sbmC* sites. A single outlier was identified encompassing *IHFb* and *sbmB* sites.

From these results, we derived a dsDNA of 170 bp termed ds_-27_+143_ in Fig. 1a, ED Figs. 2c and 3 (the nomenclature of ss, ds, and ss/ds DNAs is explained in ED Fig. 2c). This DNA encompasses the protected region and additionally the *traI_helicase_* site and the full *sbmC* site. Using ds_-27_+143_, a relaxosome complex termed “ds_-27_+143_-R” was assembled.

**Fig. 3.**
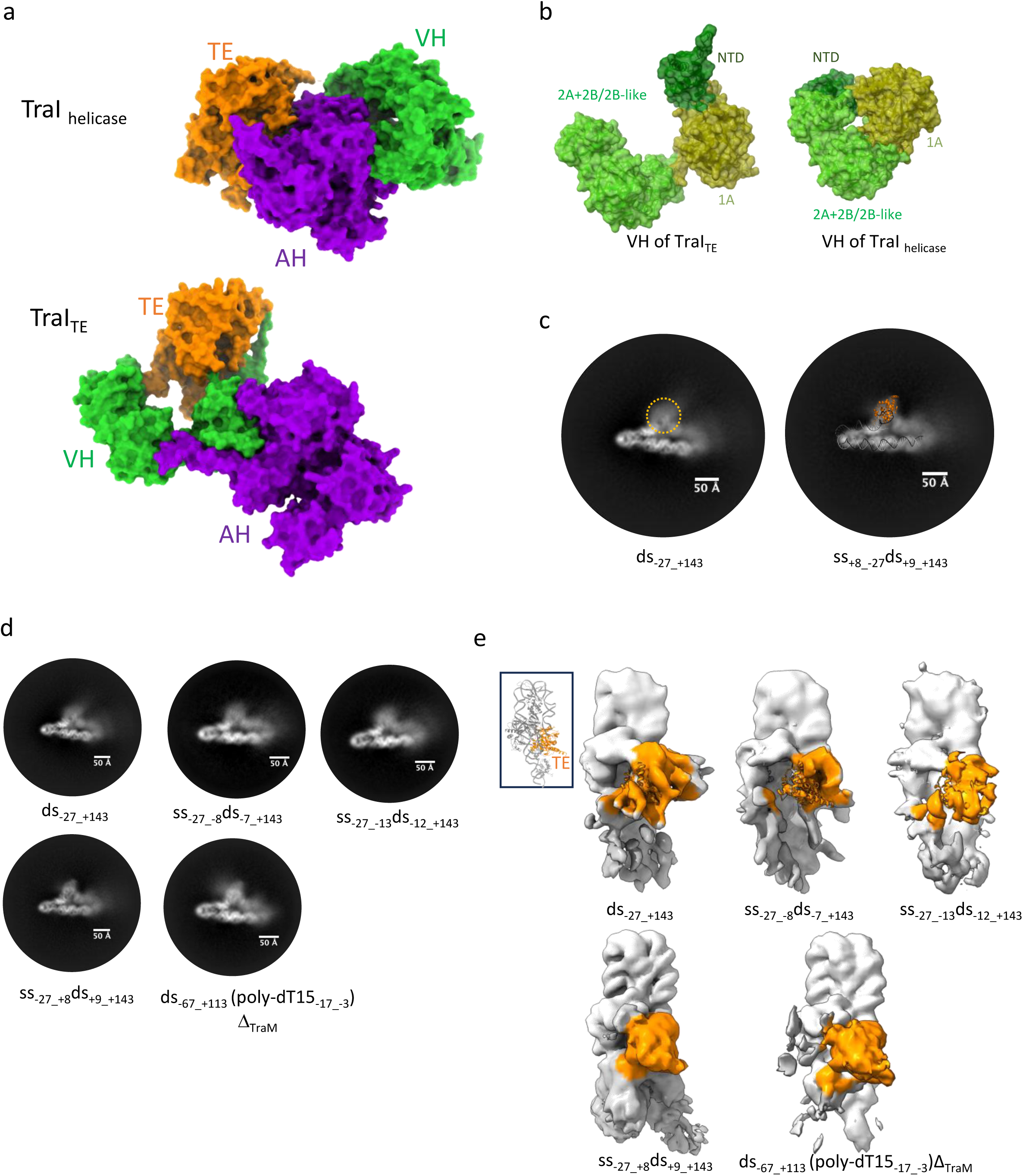
TraI conformational changes, requirement for bubble formation 3’ of *nic*. a, TraI conformational changes. TraI TE, VH and AH domains are shown in surface representation, colour-coded orange, green and violet and labelled correspondingly. Top: TraI’s helicase mode (TraI_helicase_). Bottom: TraI’s TE mode (TraI_TE_). See main text for definition of TraI_TE_ and TraI_helicase_. b, TraI VH domain in the relaxosome (TraI_TE_ at left) versus in TraI_helicase_ (right). VH_NTD_, VH_1A_ and VH_2A+2B/2B-like_ are shown in dark green, olive and green surface, respectively. In TraI_TE_, VH_NTD_ and VH_2A+2B/2B-like_ is found rotated by 47° and 95° relative to VH_1A_, respectively. c, The TE domain of ds_-27_+143_-R structure is flexibly positioned but locates in a region similar to that of the TE domain of ss_-27_+8_ds_+9_+143_-R. Two 2D-classes in the same orientation are shown, one for ds_-27_+143_-R (left) and one for ss_-27_+8_ds_+9_+143_-R. Left: a “fuzzy” density in shown in dash-lined orange circle. Right: the TE and DNA of ss_-27_+8_ds_+9_+143_-R is mapped on the 2D-class in such a way that its DNA is placed on top of the DNA’s 2D-projection. This positions the TE in the density shown at left, demonstrating that this density is generated by the ds_-27_+143_-R TE domain. However, the density is “fuzzy”, indicating the domain is flexibly located. Thus, TraI in ds_-27_+143_-R is likely similarly positioned as in TraI_TE_. d, Requirement for a DNA bubble 3’ of *nic*. Five 2D-classes are shown for five different relaxosome complexes reconstituted using five different types of DNAs. These DNAs are as described in ED Figs. 2c and 3. Two types of classes are observed, regrouped into 2 rows. The first row reports on 2D-classes where the TE region is fuzzy, i.e. in those complexes, the TE domain is not bound to its binding site. The second row reports on 2D-classes where the TE region is well defined i.e. in those complexes, the TE domain is engaged with its binding site. See main text for description and interpretation of results. e, Requirement for a DNA bubble 3’ of *nic* (continued). Same as panel d but reporting the corresponding 3D maps. The top row shows the 3D maps where the TE region is disordered: A map of similar resolution from initial processing of ds_-27_+143_-R dataset is shown for comparison. Bottom row shows the maps where TE is ordered, indicating its binding to the ssDNA. A map of similar resolution of ss_-27_+8_ds_+9_+143_-R is shown for comparison. Regions of the maps corresponding to the TE domain is coloured orange. See Methods section and ED Table 2 for details. Inset shows the relaxosome structure in ribbon in the same orientation as the maps and with the TE domain coloured in orange.

## Determination of the Cryo-EM structure of ds_-27_+143_-R

The structure of the ds_-27_+143_-R complex was derived from a “locally-refined” cryo-EM map with average resolution of 3.78 Å (ED Figs. 4, 5a,g; ED Tables 1a,b; also see definition of “locally-refined” versus “global” in ED Fig. 5). The density was of sufficient quality to build a model that includes (Fig. 1b): i- a large region of dsDNA from base pairs +12 to +94 (sequence in bold in Fig. 1a), ii- the IHF α and β chains, iii- a train of three TraY molecules and iv- two sub-domains of the VH domain of TraI, termed 2A, and 2B/2B-like (labelled VH_2A+2B/2B-like_ in Fig. 1b; see details of the TraI sub-domain structure in ED Fig. 1c,d)^17^.

**Fig. 4.**
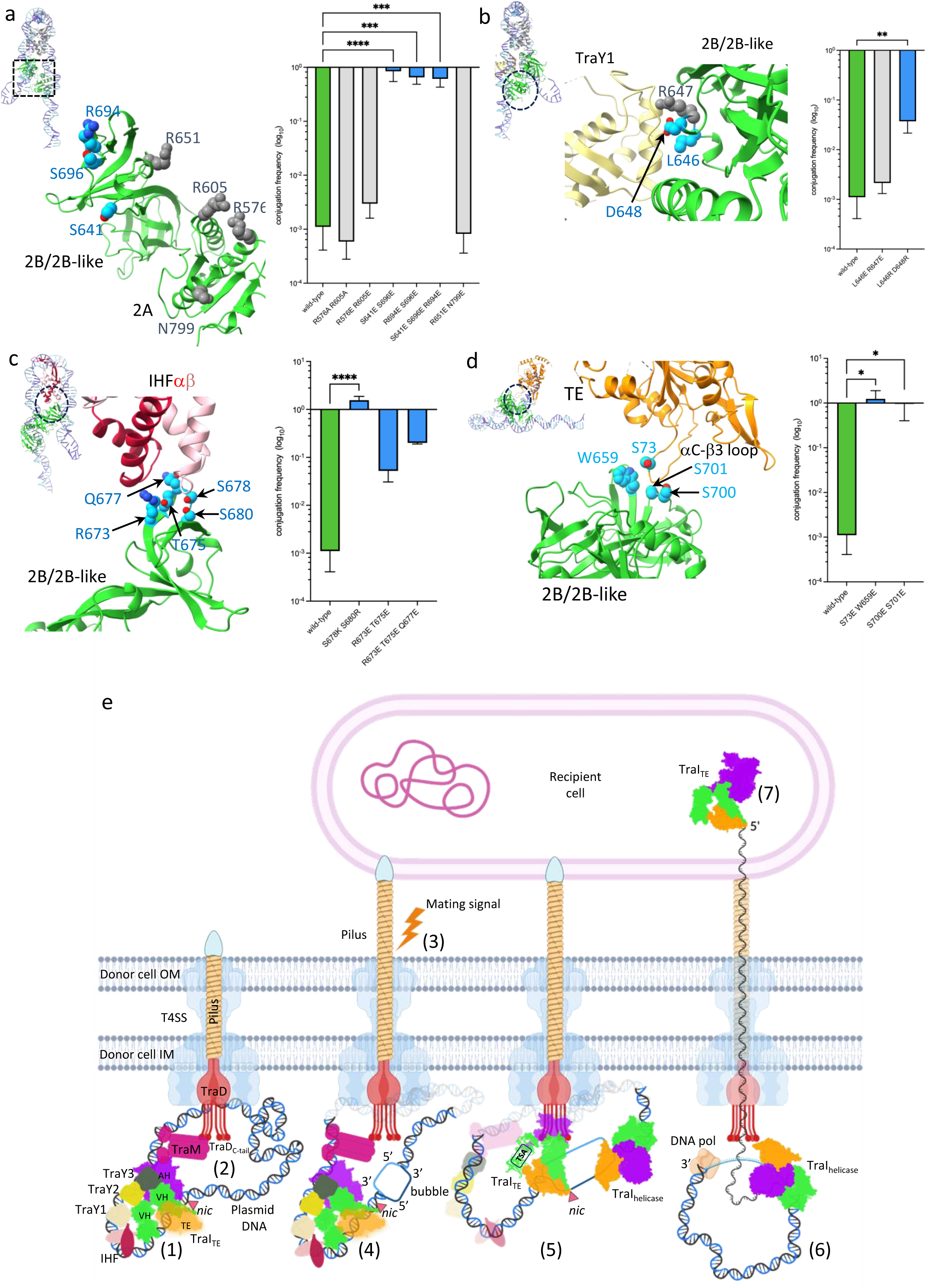
Mutational study of the relaxosome and Mechanistic model of conjugation. a-d, In each panel, the structure of the region of TraI where the mutations were introduced is shown in ribbon, with mutated residues in sphere representation colour-coded in blue or grey depending on whether they result in increased conjugation or not, respectively. Also reported is the log 10 bar graph of the conjugation rates observed for each of the mutant derivatives as described in the methods section. Colour-coding of bars is green, grey and blue for wild-type, mutants with no impact on conjugation, and mutants displaying increased conjugation, respectively. Ordinary one-way ANOVA with Holm-Šídák correction evaluated statistical significance between wild-type conjugation frequencies and complementation by mutant derivatives affecting each interface hub. n=4 for wild-type, n=3 for mutants * P < 0.05, ** P < 0.01, *** P < 0.001, **** P < 0.0001. Inset for each panel: zoom-out showing the entire relaxosome structure in the same orientation as main panel. a, TraI VH_2B/2B-like_ interaction with DNA. b, TraI VH_2B/2B-like_ interaction with TraY1. c, TraI VH_2B/2B-like_ interaction with IHF. d, TraI VH_2B/2B-like_ interaction with TraI TE. e, The molecular details of conjugation in the donor cell. Shown are the outer and inner membrane (OM and IM, respectively), the T4SS (light blue), TraD (red), the pilus (yellow) and the relaxosome proteins TraM (magenta), TraY1-3 (pale yellow, yellow and olive green, respectively), IHF heterodimer (pink and red) and TraI_TE_/TraI_helicase_ (domains color coded orange for TE, green for VH and violet for AH). The plasmid DNA is colored light blue for the R-strand and dark blue for T-strand. ‘*nic*’ site is indicated by an orange filled triangle. When the TE domain of TraI_TE_ is not engaged with its binding site, it is shown in fuzzy orange to indicate conformational flexibility, while when engaged with its binding site, it is shown unfuzzy to indicate an ordered structure. Translocation signal A (TSA) on TraI_TE_ is labelled and shown as cartoon in step (5). The steps are as described in main text.

Overall, the salient feature of this structure is an asymmetric U-shaped dsDNA hairpin, generated by IHF binding. The region of ordered dsDNA in the density stretches over 83 bp, 52 bp on one side of IHF containing the TraY-binding sites (termed the TraY-binding arm of the dsDNA hairpin in Fig. 1b; bp_+43_ to bp_+94_), and 31 bp on the other side (termed the *nic* arm of the dsDNA hairpin in Fig. 1b; bp_+12_ to bp_+42_). On the former arm, three bound TraY proteins are observed, two repeated directly (TraY1 and TraY2) and the other inverted (TraY3). This train of three TraY molecules induces longitudinal bending of the dsDNA along its long axis, changing its trajectory. On the *nic* arm, a combination of sequence-induced dsDNA bending^39^ between bp_+20_ to bp_+34_ and dsDNA bending induced by VH_2A+2B/2B-like_ binding is observed (ED Fig. 6).

The VH_2A+2B/2B-like_ module interacts with the two arms of the dsDNA hairpin and also with IHF and TraY. This dense network of interactions is seen in all relaxosome structures presented here (see details below).

## Towards reconstructing the entire relaxosome structure

We sought next to obtain a more complete relaxosome structure by experimenting with DNA sequences and ss/ds hybrid substrates. Previous X-ray crystallography of the TraI TE domain alone mapped its binding site to 11 nucleotides spanning *nic* (nucleotides -2 to +9 in Fig. 1c)^40,41^. Therefore, we designed a heteroduplex DNA termed “ss_-27_+8_ds_+9_+143_”, containing a single-stranded (ss) T-strand region from -27 to +8 and a double-stranded (ds) region from +9 to +143 (Fig. 1c, ED Figs. 2c, 3). The corresponding relaxosome complex is termed “ss_-27_+8_ds_+9_+143_-R”. This structure was built into a locally-refined cryo-EM map with an overall resolution of 3.45 Å (ED Figs. 7a, 5b; ED Tables 1a,b).

The following differences were readily apparent compared to the ds_-27_+143_-R structure (Fig. 1d): i- the TE domain of TraI was entirely observable and could be built in the density as well as the portion of the T-strand bound to it (nucleotides -2 to +9), ii- dsDNA from base pairs +10 to +93, VH_2A+2B/2B-like_, both chains of IHF, as well as the three TraY molecules present in the ds_-27_+143_-R structure were better defined in the density and improved models could therefore be derived, and iii- density for the NTD and 1A domains of VH (VH_NTD+1A_) exhibited clear secondary structural elements in which a main-chain model could be fitted in density.

Additional densities were observed in the “global” density map of the ss_- 27_+8_ds_+9_+143_-R complex (ED Fig. 5b,h). These additional densities may correspond to the AH and CTD domains of TraI or/and to TraM, a protein that is also part of the complex. To locate these domains/proteins within these densities, we solved the cryo-EM structures of the ss_-27_+8_ds_+9_+143_-R complex where i- a version of TraI lacking the AH and CTD domains was used (TraI_1-863_; complex termed ss_-27_+8_ds_+9_+143_-RΔ_AH+CTD_; ED Figs. 7b, 5c,i) or ii- TraM was omitted in complex formation (complex termed ss_- 27_+8_ds_+9_+143_-RΔ_TraM_ (ED Figs. 7c, 5d,j). Results of global map superpositions for both ss_-27_+8_ds_+9_+143_-RΔ_AH+CTD_ and ss_-27_+8_ds_+9_+143_-RΔ_TraM_ with ss_-27_+8_ds_+9_+143_-R are shown in ED Fig. 8a,b, respectively. These identified the density for TraM and for TraI AH and CTD, the models^17,19,34^ of which were subsequently fitted in the corresponding density. Further validation of TraM docking is detailed in ED Fig. 10c. Together, these results provide the basis to reconstruct a fully-assembled relaxosome (Fig. 1e).

## Structure of the fully-assembled relaxosome

The structure of the ss_-27_+8_ds_+9_+143_-R complex consists of successive but overlapping interaction centres analogous to railway hubs organised along a rigid track of DNA (Fig. 2a). Four hubs each combining protein-DNA and protein-protein interactions can be defined. Hub 1 made of IHF/TraI VH_2A+2B/2B-like_/TraY1 and dsDNA between base pairs +13 and +74 (Figs. 2b,c and ED Fig. 9a,b), Hub 2 formed by a “train” of three TraYs and dsDNA between base pairs +61 and +90 (Fig. 2d and ED Fig. 9c), Hub 3 made of TraI TE, TraI VH_2B/2B-like_, and part of the *nic* arm of the DNA hairpin that includes the ssDNA between -2 and +9 and dsDNA between +10 and +15 (Figs. 2e,f and ED Fig. 10a,b), and Hub 4 made of TraM and dsDNA between base pair +93 and +117 (ED Fig. 10c,d).

Hub 1 is the largest hub in terms of buried surface area (∼8000 Å^2^). Interactions between IHF chains and between IHFαβ and its dsDNA-binding site (between bp+23 and bp+60; Fig. 2b) account for much of it (5300 Å^2^). The 160° bending induced by IHF binding is caused by the intercalation of two β-hairpins (β3 and β4) within the DNA’s minor groove (Fig. 2b) ^22^. In the hub, IHFα reaches out to TraY1 (158 Å^2^) while both its chains interact with the TraI VH_2B/2B-like_ sub-domain (∼400 Å^2^). Furthermore, TraY1 provides a binding interface to TraI VH_2B/2B-like_ (410 Å^2^) (Fig. 2b). These interaction areas are not individually extensive but their cumulative effect on TraI VH serves to calibrate TraI VH_2A_ domain against its dsDNA binding site on the TraY arm of the dsDNA hairpin between bp +61 and +74 (Fig. 2c, ED Fig. 9b).

Hub 2, composed of the train of TraYs, is characterised by a large network of protein-protein interactions between TraY subunits and protein-DNA interactions that anchor the train to 3 binding sites on the dsDNA (Fig. 2d, ED Fig. 9c). TraY is an RHH (ribbon-helix-helix) family protein, which are usually dimers. In contrast, F TraY is a monomer containing all the secondary structures and tertiary fold of the RHH dimer in one single polypeptide. Overall, the three TraY molecules bury 2833 Å^2^ of surface area upon interaction with their binding sites (see details in ED Fig. 9c). Interaction between TraY molecules is also extensive. Remarkably, TraY appears to be able to use very different surfaces to mediate protein-protein contacts (details in ED Fig. 9c).

In Hub 3, TraI TE is observed bound to its previously characterised ssDNA-binding site between -2 and +9^41^. However, in marked contrast with previous observations (ED Fig. 10a,b), it is also seen to make extensive contacts with the dsDNA between +10 and +15 (Fig. 2e). The latter interface accounts for nearly half of the total TE-DNA interaction area in the relaxosome (1295 Å^2^ TE-dsDNA versus 1365 Å^2^ for TE-ssDNA interactions). Interactions details are described in Fig. 2e and ED Fig. 10a,b. Interestingly, residues in the β9-αF loop (Fig. 2e) displaces the R-strand base (a C) of the CG +9 base pair out of its pairing arrangement, resulting in only G_+9_ of the T-strand being visible in the electron density. Thus, the TraI TE domain is able to melt/unwind the +9 base pair, an observation that prompted us to ask whether this ability to melt/unwind dsDNA extends beyond +9 (see below). Finally, the TE domain provides a small but stabilising interface that anchors TraI VH_2B/2B-like_ sub-domain within interactions with the dsDNA near the ss/ds DNA junction (Fig. 2f).

Next, to assess the magnitude of the conformational change needed in the T-strand to go from unbound to TE-bound, we *in silico* extended the ds_-27_+143_ DNA to include *nic* and superimposed it onto the ss/dsDNA of the ss_-27_+8_ds_+9_+143_-R structure (ED Fig. 10e). We observe that not only has the *nic* site moved 29 Å away from its unbound location, but also that the T-strand on TE-binding has undergone a U-turn, imposed by its interaction with TE and notably residues in the β10-β11 region (Fig. 2e). Thus, *oriT* is subjected to two topological U-turns, one in its dsDNA region upon IHF-binding, and another one in its ssDNA region upon TE-binding.

TraM is made of two domains (ED Fig. 1f): an N-terminal domain (NTD) that interacts with DNA and a C-terminal domain (CTD) responsible for tetramerization and known to interact with the coupling protein TraD^34,42^. Thus, TraM can also be defined as a “hub” of protein-DNA and protein-protein interactions. This hub, Hub 4 (described in ED Fig. 10c,d), is distinct from hubs 1-3 in that it not only mediates interactions between intra-relaxosome components but also with the T4SS^42^.

A striking feature of the ss_-27_+8_ds_+9_+143_-R structure is the very large DNA footprint made by the various proteins involved (Fig. 2g). Protein-DNA interactions involve a staggering 9022 Å^2^ of surface area, with IHF and TraI TE accounting for more than half of this total. When the TraM/DNA interaction is added, over 10,000 Å^2^ of buried surface area are involved in assembling the relaxosome components on *oriT*.

## The TE and Helicase binding modes of TraI

With the ss_-27_+8_ds_+9_+143_-R relaxosome structure, we capture a state of TraI where the protein engages with its TE-binding site, revealing its TE mode, TraI_TE_. In this conformational state, TraI has undergone a dramatic re-arrangement of its domain and subdomain structure compared to its helicase mode (Fig. 3a,b): i- while the AH domain was proximal to the TE domain in TraI_Helicase_, in TraI_TE_, it is the VH domain that is now proximal to that domain (Fig. 3a); ii- In TraI_TE_, the VH domain itself has undergone a large re-arrangement of its sub-domains, with the NTD and the 2A+2B/2B-like module pivoting 47° and 95° degrees each relative to the 1A sub-domain (Fig. 3b). The AH and CTD domain of TraI remain largely semi-unstructured, lying in a region of the relaxosome that we were able to locate, consistent with the observation that TraI_TE_ is sensitive to mild proteolysis i.e. is semi-flexible. Apparently, this region makes no specific and stabilising contacts with proteins or DNA nearby. Finally, when a model of the ss_-27_+8_ds_+9_+143_-R is placed onto the 2D-classes of ds_-27_+143_-R, we observe that the TE domain maps to a region of the 2D-class that forms a fuzzy protrusion from the dsDNA (Fig. 3c), indicating that this region of ds_-27_+143_-R, which has poor density and could not be interpreted, corresponds to TraI TE. Thus, when in a dsDNA-bound relaxosome, TraI is in its TraI_TE_ mode.

## TE binding requires the formation of a bubble at the helicase binding site

As mentioned above, in the ss_-27_+8_ds_+9_+143_-R structure, the R-strand base of the first double-stranded pair of the ss/dsDNA duplex (pair +9 in Fig. 2e) is not observed in the density due to its displacement by the β9-αF loop. We asked whether TraI TE can melt a more extended region of dsDNA.

Thus, we reconstituted relaxosome complexes with decreasing ssDNA length, progressively extending the dsDNA part by increments of 5 nucleotides towards and beyond the *nic* site (ED Figs. 2c and 3). The DNA employed were: ss_-27_+3_ds_+4_+143_, ss_-27_-3_ds_-2_+143_, ss_-27_-8_ds_-7_+143_ and ss_-27_-13_ds_-12_+143_. Cryo-EM datasets were collected for each complex. 2D classes and 3D maps were computed (see Methods and ED Table 2). The ss_-27_+3_ds_+4_+143_-R and ss_-27_-3_ds_-2_+143_-R were solved to high resolution and are virtually identical to ss_-27_+8_ds_+9_+143_-R with DNA observed bound to a well- ordered TraI TE domain (ED Fig. 5e,f, ED Tables 1a,b). For ss_-27_-8_ds_-7_+143_-R and ss_- 27_-13_ds_-12_+143_-R, only 2D classes and initial low resolution 3D maps were obtained because it became clear early on in the processing that their structure was similar to that of ds_-27_+143_-R, i.e. with a disordered TraI TE domain (Figs. 3d,e). Thus, in the ss_- 27_+3_ds_+4_+143_-R and ss_-27_-3_ds_-2_+143_-R complexes, the TraI TE domain engages with its binding site while, in the ss_-27_-8_ds_-7_+143_-R and ss_-27_-13_ds_-12_+143_-R complexes, it does not. Together, these results indicated that TraI TE is unable to melt sequences beyond base pair -3. A corollary conclusion is that a single-strand of DNA is required before base pair -3 in order for TraI TE to melt and engage its binding site. Accordingly, we asked whether a bubble immediately 3’ to the *nic* site would be sufficient for TraI TE to melt its DNA-binding site. To test this hypothesis, a dsDNA containing 15 bases (poly dT15) was inserted between position -17 and -3 as described in ED Figs. 2c or 3. The DNA was extended on the 3’ side of the T-strand to stabilize the bubble. TraM and its binding site were not included. This DNA is referred to as ds_-67_+113_(poly-dT15_- 17_-3_). A limited cryo-EM dataset as above was collected, 2D classes and a 3D map were obtained and shown to resemble those of ss_-27_+8_ds_+9_+143_ indicating an ordered TE domain (Figs. 3d,e). Thus, a single strand bubble immediately 3’ of *nic* is required for the binding reaction to occur.

## Validation of the relaxosome structure

To validate the structures presented here, we initiated an extensive mutational study, targeting residues in three hubs, Hub1-3 (Fig. 4a-d and ED Table 3). All mutations were made in *traI* and analyzed for conjugative plasmid transfer activity. Firstly, we targeted residues interacting with dsDNA (Fig. 4a). Both interactions of VH_2B/2B-like_ residues with the *nic* arm of the dsDNA hairpin and of VH_2A_ residues with the TraY arm were tested by generating negatively charged substitutions. Double and triple mutants were made (Fig. 4a). We observed either no effect for residues of VH_2A_ or very large increases in conjugation rates for residues in VH_2B/2B-like_ (nearly 300-fold for S641E- S696E for example), pointing to an interaction hub that plays important roles. Very similar results are obtained when mutating some residues at the interface between TraI VH_2B/2B-like_ and TraY1 (Fig. 4b), between TraI VH_2B/2B-like_ and IHF (Fig. 4c), and between TraI VH_2B/2B-like_ and TraI TE (Fig. 4d). These results all emphasize the functional importance of the various hubs we have characterised in the structure. But perhaps more significantly, they point to the possibility that our structure represents a quiescent or pre-initiation state of the relaxosome, the activation of which can be triggered readily by changes in the structure, possibly involving disruption of the key hub interactions analysed here. The activation step is predicted to occur in response to signals such as pilus termination or the presence of a recipient cell^43,44^. Our mutational study results indicate that changing any combination of residues disrupt a finely-tuned quiescent complex that can easily be triggered by extracellular signals.

## Mechanism of conjugation

In this report, we present the structure of the relaxosome in various DNA-bound states. From the mechanistic insights obtained here, a molecular-level mechanistic model for relaxosome recruitment and activation emerges (Fig. 4e). It starts with the formation of a quiescent relaxosome at *oriT* with TraI in its TraI_TE_ mode (step 1) and its subsequent recruitment to the T4SS (step 2) via the interactions of the coupling protein TraD with TraM and of TraI TSA and TSB modules with an unknown receptor^45,46^. Once pilus biogenesis is completed and a recipient cell has contacted a donor cell, a signal is sent (step 3) that activates the relaxosome by generating an ssDNA bubble’ immediately 3’ to the *nic* site on the T-strand (step 4). This allows i- the TE of TraI_TE_ to access its binding site as in the structure of ss_-27_+8_ds_+9_+143_-R and ii- the loading of TraI_helicase_^17^ (step 5). The former functions in its so-called TraI_TE_ mode when bound to *traI_TE_* site (ED Fig. 1b), proceeds to cleave *nic* and attach covalently to the 5’phosphate resulting from the nicking reaction, while the latter when bound 3’ of *nic* (*traI_Helicase_* site in ED Fig. 1b) initiates DNA unwinding (step 6). In a final step, the DNA is driven through the T4SS, through the pilus, and finally injected into the recipient cell (step 7).

## Data availability

EM maps and atomic models were deposited to the Electron Microscopy Data Bank (EMDB) and Protein Data Bank (PDB) databases. Accession codes can be found in ED Tables 1 a, b and 2. All constructs (wild type and mutants) used in this study can be obtained on request to GW.

## Acknowledgements

We would like to thank Dr. David Houldershaw for IT support and Drs. Natasha Lukoyanova and Shu Chen for collection of data on the in-house Krios. This work was supported by Wellcome grants 098302 and 217089 to GW and the Austrian Science Fund FWF Doc50 Molecular Metabolism to ELZ. Cryo-EM data were collected at the ISMB EM facility at Birkbeck College, University of London with financial support from Wellcome Trust (202679/Z/16/Z and 206166/Z/17/Z) and at the UK national electron Bio-Imaging Centre (eBIC).

## Author contributions

SMW purified the DNA and protein components, established the conditions for relaxosome assembly and set up the DNaseI footprinting protocol. SMW conducted NS and cryoEM sample preparations, data collection, processing, model building and refinement. SMW and GW carried out structural analyses. SR, SK, and ELZ performed the mutational analysis of *traI*. AI was involved with relaxosome characterisation in the initial phase of the project. SMW, ELZ and GW wrote the manuscript with inputs from SR. GW supervised the work.

## Methods

### Cloning, protein expression and purification

All strains, plasmids, constructs, and primer sequences used in this study are listed in ED Tables 4a-d, respectively. A tagless construct of IHF was generated by PCR amplification excluding the tag from a vector where *ihfA* and *ihfB* genes were cloned in tandem in the MCS of pRSF-1b plasmid (Novagen), to generate pRSF-IHF. The *traM* and *traY* genes synthesised and cloned into the NcoI and HindIII sites of pET28a vector were purchased from GenScript. All TraI expression constructs used in this study were engineered from pTrc99a-TraI^48^ harbouring wild-type *traI* gene cloned into the MCS. Mutagenesis following the In-Fusion cloning protocol was used to generate the single Y16F mutant TraI variant (TraI_Y16F_). Expression construct for TraI_Y16F_ domain deletion variant TraI_ΔAH+CTD_ was generated by PCR amplification of pTrc99a-TraI, excluding the region of TraI between residue 835 to the C-terminus and annealing by In-Fusion seamless cloning technology (Takara Bio). All constructs were verified by DNA sequencing (Eurofins Scientific).

IHF was purified as a heterodimer with a protocol adapted from Nash *et al.*^49^. Briefly, *E. coli* BL21 (DE3) cells (ThermoFisher) transformed with pRSF-IHF were grown in 2L of LB media supplemented with 50 µg/ml kanamycin at 37°C. Upon reaching an OD_600_ of 0.7, expression was induced with IPTG at a final concentration of 1mM and incubated overnight at 18°C. The cells were harvested by centrifugation, resuspended in 20mM Tris-HCl pH 7.5, 100 mM NaCl (buffer A), one protease inhibitor cocktail tablet (Roche) and lysed by sonication (10-15 cycles of 5 s ON and 10 s OFF using a Sonics Vibra Cell VCX 130 sonicator set at 30% amplitude). The lysed cells were centrifuged, and the resultant supernatant was subjected to stepwise precipitation with ammonium sulphate to 50% and 70% saturation. The 70% precipitate was dialysed in 20mM Tris-HCl pH 7.5, 50 mM NaCl (buffer B) and applied to a 5 ml HiTrap Heparin column (GE Healthcare) equilibrated with the same buffer. A gradient elution was performed using 50mM-1M NaCl in 20mM Tris-HCl pH7.5, with fractions containing IHF heterodimer then concentrated and subjected to size exclusion chromatography using a Superdex 75 Increase 10/300 column (GE Healthcare) with buffer A as the mobile phase. The fractions containing the protein were concentrated, flash frozen in liquid nitrogen and stored in -80°C.

TraM was purified as follows. C41 (DE3) cells (Lucigen) harbouring pET28a- TraM plasmid were grown in 1L LB media with 50 µg/ml kanamycin at 37°C. On reaching an OD_600_ of 0.7, protein was induced by IPTG at a final concentration of 1mM and incubated further at 18°C, overnight. The pelleted cells were resuspended in buffer A with a protease inhibitor tablet (Roche). Lysis was carried out by sonication as described for IHF and the clarified lysate was subjected to precipitation with ammonium sulphate to 40% and 100% saturation. The 100% pellet was dialysed in buffer B and applied to a 5 ml HiTrap Heparin column (GE Healthcare). Elution was carried out using a gradient of 50mM-1M NaCl in 20mM Tris-HCl pH7.5 with fractions containing TraM, detected by SDS-PAGE, pooled and diluted until the conductivity was equivalent to buffer B. This was applied to a 6 ml resource Q column (Cytiva) equilibrated with buffer B. A linear gradient of 50mM-1M NaCl resulted in the elution of proteins and the desired fractions were further diluted with buffer B. This was applied to a 5 ml HiTrap Blue column (Cytiva) and gradient elution was carried out with 50mM- 2M NaCl. The fractions having the protein were then concentrated and subjected to size exclusion chromatography using a Superdex 200 Increase 10/300 column (GE Healthcare) equilibrated with buffer A. Fractions containing purified TraM tetramers were concentrated, flash frozen in liquid nitrogen and stored in -80°C.

TraY was expressed in C41 (DE3) cells transformed with pET28a-TraY and lysed by sonication following the steps described for TraM. The clarified lysate was then loaded directly onto a 5 ml HiTrap Q column (Cytiva) connected in tandem to a 6 ml resource S column (Cytiva) such that the flow-through from the Q column passes through resource S. The Q column was then disconnected and gradient elution with 100mM-1M NaCl in 20mM Tris-HCl pH7.5 was carried out on the resource S column. The fractions containing TraY were concentrated and further fractionated on a Superdex 75 Increase gel filtration column equilibrated with buffer A. The purified fractions were further concentrated and stored in -80°C after flash-freezing in liquid nitrogen.

TraI, TraI_Y16F_ and its truncation variant TraI_ΔAH+CTD_ were expressed in BL21 (DE3) cells (ThermoFisher) harbouring the corresponding pTrc99a clone, grown in 2L of LB media supplemented with 100 µg/ml carbenicillin at 37°C. Once the cells reached an OD_600_ of 0.7, expression was induced with a final IPTG concentration of 1 mM and incubated overnight at 18°C. The cells were harvested by centrifugation, resuspended in buffer A with one protease inhibitor tablet (Roche) and lysed by sonication as described above. The clarified cell lysate was subjected to ammonium sulphate precipitation (0.3g/ml) with the resulting pellet dialysed in buffer B after resuspension in the same buffer. The solution after dialysis was applied onto a 5 ml HiTrap Heparin column (GE Healthcare) equilibrated with the same buffer. A linear gradient elution was performed using 50mM-1M NaCl in 20mM Tris-HCl pH7.5 and the eluted fractions containing the protein diluted until conductivity was equal to buffer B. This was then applied to a 5 ml HiTrap Q column (GE Healthcare) equilibrated with buffer B. Gradient elution was carried out using 50mM-1M NaCl in 20mM Tris-HCl pH7.5 and fractions having the protein were concentrated and subjected to size exclusion chromatography on a Superose 6 Increase column equilibrated with buffer A. The purified protein fractions were concentrated and stored in -80°C after flash-freezing in liquid nitrogen.

### DNA synthesis and purification

Synthesis of duplex *oriT_316_* DNA fragment was performed by large scale PCR amplification of the *oriT* region of F plasmid synthesised and cloned in pUC57 (pUC57- *oriT_316_*), purchased from Genscript. A 5 ml PCR reaction containing 5 μg of template DNA, 0.5 μM final concentrations each of primers (ForiT_F, ForiT_R), 2X master mix of CloneAmp HiFi PCR Premix (Clonetech, USA) was carried out in a thermocycler with the following PCR conditions: 98 °C for 2 min, and 30 cycles of 98 °C for 10 s, 55 °C for 5 s, 72 °C for 5 s , and a final extension at 72 °C for 10 min. The DNA was precipitated by addition of 1/10 volume of 3M sodium acetate pH 5.2, 3 times volume of 100% ethanol and incubated o/n in -80 °C. Precipitated DNA was recovered by high-speed centrifugation, after which the supernatant was discarded. The pellet was dried at RT, washed with 100% isopropanol, further dried at RT and resuspended in 1 ml buffer A and a further step of purification was carried out on a Superose 6 Increase column equilibrated with buffer A. Fractions were assessed on a 4-20% TBE PAGE gel (ThermoFisher) after which the pure fractions were pooled giving a yield of 60-80 μg DNA, and stored in -20°C.

For generation of the ds_-27_+143_ DNA by restriction digestion, a sequence containing five consecutive ds_-27_+143_ DNA fragments separated by six HindIII restriction sites, synthesised and cloned in pUC57-mini vector (pUC57mini- ds_- 27_+143__5X), was purchased from Genscript. A large-scale restriction enzyme reaction mix containing 500 μg of plasmid, 10,000 units of HindIII-HF (NEB) and 1X CutSmart buffer was made up to a final volume of 12.5 ml with MilliQ water. The reaction was incubated at 37°C for 1 hour followed by heat inactivation at 80°C for 20 mins and any aggregates were removed by centrifugation. The 12.5 ml mixture was concentrated to a volume of 100 μl in an Amicon Centricon Filter (Millipore) with a 3 kDa cut-off. The digested DNA was size-fractionated on a native-PAGE (5% TBE gel, 40% acrylamide/bis-acrylamide) using Mini Prep Cell (Bio-Rad). The fractions were inspected on a 4-20% TBE PAGE gel (ThermoFisher) and the ones containing ds_- 27_+143_ DNA were desalted with buffer A and concentrated, in a 3 kDa cut-off Amicon Centricon Filter (Millipore), giving a yield of 50-60 μg DNA, and stored in -20°C.

All the other DNA constructs used for relaxosome reconstitution were generated from PAGE purified Ultramer^TM^ Oligos purchased from IDT. The corresponding R- and T- strand oligos (ED Table 4e) were mixed at 1:1 molar ratio at a final concentration of 10 μM in a volume of 100 μl and annealed by incubating at 98°C for 5 min, lowering the temperature at a rate of 1°C/min until reaching 24°C, in Annealing Buffer (50 mM TrisHCl pH7.2, 50 mM NaCl, 1 mM EDTA, 1 mM MgCl_2_). The annealed DNA was purified by electrophoresis though a native acrylamide gel column (12% TBE gel, 40% acrylamide/bis-acrylamide) using Mini Prep Cell (Bio-Rad). The fractions were analysed for purity on a 4-20% TBE PAGE gel (ThermoFisher) and the pure fractions were desalted with buffer A and concentrated, in a 3 kDa cut-off Amicon Centricon Filter (Millipore), yielding 60-80 μg of DNA. Correct annealing of all annealed DNA heteroduplexes were verified by assessing the length of the two constituent ssDNA strands using a 10% TBE-Urea denaturing gel (ThermoFisher). The ds_- 67_+113_(poly-dT15_-17_-3_) was additionally verified to confirm the unpaired region by S1 nuclease (Thermofisher) digestion to release its release right and left duplexes of 115 and 50 bp, respectively. The purified oligos were stored in -20°C.

### Relaxosome DNaseI footprinting and sequencing of resulting fragments

The relaxosome complex was formed by mixing 0.25 μM *oriT_316_* DNA, molar excess of proteins at 8 μM each of IHF and TraM, 10 μM TraY, 5 μM TraI, 100 μl of 10X DNase I buffer (Roche) made up to a volume of 1 ml with Buffer A and was allowed to incubate for 30 min at RT. To this, 200 units of DNase I recombinant RNase-free (Roche) was added and further incubated for another 45 min at RT. Following this, an equal volume of premixed phenol:chloroform:isoamyl alcohol 25:24:1 (v/v) (ThermoFisher) was added, sample was mixed vigorously and centrifuged at 17,000 x g for 10 min. The upper aqueous phase containing DNA was transferred to a new tube, mixed with 1/10 volume of 3M sodium acetate pH 5.2, 3 times volume of 100% ethanol and incubated o/n in -80°C. Precipitated DNA was recovered by centrifugation at 17,000 x g for 30 min at 4°C, after which the supernatant was discarded and the pellet was washed with 1 ml Isopropanol. The sample was centrifuged at 17,000 x g for 30 min at 4°C, supernatant discarded and the pellet was dried at RT. The air-dried pellet was resuspended in 100 μl MilliQ and purified by 1.5% agarose gel electrophoresis followed by extraction of DNA fragments from the excised gel band with Nucleospin gel and PCR Clean-up kit (Takara).

To facilitate cloning of DNA fragments into pCR-BluntII-TOPO vector (ThermoFisher), the fragments were subjected to blunting and dephosphorylation. For the blunting reaction 2 x 20 μl reactions each containing 1 μg DNA, 100 μM of each dNTP (NEB), 1 Unit T4 DNA polymerase (NEB), 1X 2.1 Buffer (NEB) were incubated at 12°C for 15 mins followed by heat inactivation at 75°C for 20 mins. The reaction was purified with Nucleospin gel and PCR Clean-up kit (Takara) following the manufacturer’s instructions for PCR clean-up. This was followed by dephosphorylation of the blunt ends, where 8 x 20 μl reaction volumes each containing 1 pmol DNA ends, 1 X CutSmart buffer (NEB) and 1 unit of Shrimp Alkaline Phosphatase (rSAP) (NEB) was incubated at 37°C for 30 mins followed by heat inactivation at 65°C for 5 mins. The reactions were pooled together and purified with Nucleospin gel and PCR Clean- up kit (Takara) according to the manufacturer’s instructions for PCR clean-up.

A footprint library was generated by cloning the modified fragments into a pCR-BluntII-TOPO vector following instructions in the Zero Blunt TOPO PCR cloning kit (Invitrogen). Briefly, 6 μl reactions where DNA was mixed to achieve 1:10 or 1:20 molar vector-to-insert ratio, with 1 μl salt solution provided with the kit, were incubated at RT for 5 min. 4 μl of the reaction was used to transform Top10 competent cells (Invitrogen), plasmids were isolated from resultant colonies and the insert sequence established by DNA sequencing (Eurofins Scientific) (ED Table 4f).

### Relaxosome complex formation, cross-linking and purification for cryoEM

A nickase inactive version of TraI harbouring Y16F mutation (TraI_Y16F_) was used for all cryoEM complex formations^50^. For cryoEM studies, complex formation was carried out as follows. To 0.25 μM *oriT* DNA supplemented with 20 mM MgCl_2_, 20 times molar excess of purified proteins were added in the order of IHF, TraM and TraY followed by 5 times molar excess of TraI_Y16F_ or TraI_ΔAH+CTD_ in a volume of 1 ml in Buffer A. This mixture was allowed to incubate for 30 min at RT following which the unbound proteins were separated from the DNA-protein complex on a Superose 6 Increase 10/300 column (GE Healthcare), equilibrated with 20 mM Hepes pH 7.5, 100 mM NaCl (Buffer C). The peak fractions containing the relaxosome were pooled (1 ml total volume) to which Glutaraldehyde (Grade I, 25% in H_2_O, Sigma) was added to a final concentration of 0.5%. The cross-linking was allowed to proceed for 5 mins at RT and was terminated by the addition of TrisHCl pH 7.5 to a final concentration of 100 mM. The sample was subjected to a further round of gel filtration on a Superose 6 Increase 10/300 column (GE Healthcare), equilibrated with Buffer C, the fractions were pooled and concentrated using 30 kDa MWCO Microcon centrifugal filters (Millipore). The sample was analysed on a 4-12% Bolt Bis-Tris Plus gel (Invitrogen) and the gel band corresponding to the cross-linked relaxosome were excised and constituent proteins identified by peptide mass fingerprinting where appropriate. The *oriT* DNA and protein components indicated in ED Table 4g were used to form the different relaxosome complexes described in this study, but the protocol is as outlined above for each of the complexes.

### Negative stain EM

To check the quality and concentration of samples, 4 μl of 0.05 mg/ml cross-linked relaxosome was applied on glow discharged formvar-coated copper grids (Agar Scientific), incubated at RT for 1 min. The excess sample was blotted, and the grid stained with 4 μl of 2% uranyl acetate for 30 s. Excess stain was removed by blotting and the grids were allowed to dry at RT. The grids were imaged on a Tecnai T10 transmission electron microscope (Thermo Fisher Scientific) operated at 100 keV.

### CryoEM Grid preparation and data collection

For cryoEM analysis, 3 μl of the sample was applied to UltrAuFoil grids (Quantifoil, Germany; 2/2 200 mesh) that had been previously negatively glow discharged using PELCO Easiglow (Ted Pella, USA) and coated with a layer of graphene oxide^51^. The grids were incubated after sample application for 30 s in the chamber of a Vitrobot Mark IV (Thermo Fisher Scientific, USA) at 4 °C and 95% humidity and excess sample was blotted and vitrified in liquid ethane. The data were collected either at the ISMB Birkbeck EM facility or at the eBIC National facility using a Titan Krios microscope (ThermoFisher, USA) operated at 300 keV. The Krios was equipped with a post-GIF direct electron detector (Gatan,USA) and a BioQuantum energy filter (Gatan, USA) with a slit width of 20eV. Images were collected automatically using the EPU software (ThermoFisher, USA) in super resolution mode with the total dose fractionated over 50 frames. The data collection parameters used for the different datasets are listed in ED Tables 1a and 2.

### Image processing

The movies were first corrected for drift and dose-weighted using MotionCor2^52^, followed by contrast transfer function (CTF) estimation using CTFFIND v4.1^53^ within RELION^54^. Micrographs with a CTF-fit score better than 8 Å were selected for further processing. Workflows for image processing are reported in ED Figs. 4 and 7.

#### Image processing of ds_-27_+143_-R

A total of 55,914 movies were collected for ds_-27_+143_-R from three different sessions giving three datasets which were processed separately and combined at a later stage. Initially, a subset of 4000 images were selected from dataset 1, to generate an initial model to test data quality and create templates for particle picking. From the 4000 images, 953,598 particles picked by reference-free auto-picking (Laplacian-of-Gaussian) in RELION were extracted with a box size of 400 pixels (1.06 Å/pixel sampling) and subjected to several rounds of 2D classification in cryoSPARC^55^. The resulting 374,358 particles were used for *ab initio* 3D classification followed by 3D refinement (cryoSPARC) of the best class (based on discernible features) to yield an initial 3D map (cryoSPARC). The map low pass filtered to 20 Å was used to obtain 30 randomized 2D projections, which were used as templates for reference-based Gautomatch v0.56^56^ particle picking for datasets 1-3.

A total of approximately 3.5, 6 and 10 million particles were picked with permissive picking parameters for datasets 1, 2 and 3, respectively. Particles in datasets 2 and 3 were further split into smaller subsets for downstream classification steps to speed up processing. Several rounds of 2D classification in cryoSPARC resulted in a few high resolution 2D classes in a few preferred orientations but showing clear secondary structure features, and a majority of low-resolution classes comprising broken and junk particles. Selecting only the high-resolution 2D classes resulted in maps with resolution anisotropy. Therefore, to avoid throwing away rare views the selection was kept liberal at the 2D classification stage, retaining 2D classes with any semblance of particle-like features and discarding only the ‘obvious’ junk classes. Instead, to tackle heterogeneity and clean up junk, we subjected the particles from 2D classification to rigorous rounds of *ab initio* 3D classification.

Best particles were selected from successive runs of *ab initio* classification where in each round the best 3D class was selected for the next round until an *ab initio* model with high resolution features was obtained. The outcomes of datasets 1 and 2 were combined (556,418 particles) followed by 3D refinement in cryoSPARC resulting in a map of 5.97 Å. The map was used as an initial model for a round of alignment-free 3D classification in RELION with 6 classes and a regularisation parameter Tau of 16, to resolve further heterogeneity. Particles from the best two classes were combined with particles established from a similar workflow from dataset 3 and was refined to 4.31 Å resolution in cryoSPARC.

This consensus map of 337,238 particles showed disordered regions which disappeared when applying higher contour levels, indicating conformational heterogeneity. The density in the ordered region showing secondary structure details was subjected to local refinement in cryoSPARC with a mask excluding the disordered regions, yielding a map of 3.78 Å resolution. This map was used to build and refine models for *oriT* DNA, IHF, three molecules of TraY and TraI VH_2A+2B/2Blike_. We could identify a poorly resolved region suggestive of 1A of VH domain, but this region lacked secondary structure features and therefore no model was derived from it. The density in the disordered regions did not improve with further processing such as focused classification/refinement, particle subtraction etc.

#### Image processing of ss_-27_+8_ds_+9_+143_-R, ss_-27_+8_ds_+9_+143_-RΔ_TraM_, ss_-27_+8_ds_+9_+143_- RΔ_AH+CTD_

The overall strategy for the image processing of datasets with the ss/ds *oriT* heteroduplexes follows the one described above (ED Fig. 7a-c). We were able to achieve high resolution maps with fewer images as compared to ds_-27_+143_-R, since the samples seemed to have a higher degree of stability and homogeneity, likely due the binding of TraI TE which resulted in the arms of the DNA hairpin being less flexible. In the datasets of ss_-27*_+8*_ds_+9_+143_-R and ss_-27*_+8*_ds_+9_+143_-RΔ_TraM_ which were collected from regions of thicker ice, more particles in diverse orientations were observed as evidenced by 2D classes and the angular distribution calculated from cryoSPARC heat maps (ED Fig. 5b,d), yielding better 3D reconstructions in both cases.

For ss_-27*_+8*_ds_+9_+143_R, a final set of 155,077 best particles were selected, resulting in a global refined map of 3.77 Å resolution. The ordered regions showing secondary structure details in the global map were further subjected to local refinement yielding a map of 3.45 Å resolution. This locally-refined map was then used to build and refine models of the TraI TE domain bound to ssDNA, in addition to *oriT* DNA, IHF, three molecules of TraY and TraI VH_2A+2B/2Blike_. Upon inspecting the global map, secondary structural features were clearly recognizable for the NTD and 1A subdomains of VH and was used to dock AlphaFold 2 models for the same (see below).

In the global map of ss_-27*_+8*_ds_+9_+143_R we could also identify two poorly resolved regions, one next to the TraI VH domain and another at the distal end of the TraY arm of the *oriT* DNA. By comparing the global map from ss_-27*_+8*_ds_+9_+143_-RΔ_AH+CTD_ (3.93 Å, 165,577 particles; ED Fig. 5i) to the ss_-27*_+8*_ds_+9_+143_R map, we could assign this disordered region to the active helicase (AH) and C-terminal domains (CTD) of TraI. Similarly, the global ss_-27*_+8*_ds_+9_+143_-RΔ_TraM_ map (3.11 Å, 330,708 particles; ED Fig. 5j) was used to assign the density in the distal end of the *oriT* DNA to TraM and its DNA binding region.

#### Image processing of ss_-27_+3_ds_+4_+143_-R and ss_-27_-3_ds_-2_+143_-R

To further explore if the relaxosome could lead to localized melting of *oriT* near the *nic* site to facilitate TE docking, we extended the 5’ end of the R-strand by five and ten nucleotides to yield ss_-27_+3_ds_+4_+143_-R and ss_-27_-3_ds_-2_+143_-R complexes, respectively. The data collection and processing were essentially similar to that of the previous ssds constructs with the final locally refined map resolved to 3.68 Å (159,080 particles; ED Fig. 5e) for ss_-27_+3_ds_+4_+143_-R and 3.42 Å (177,743 particles; ED Fig. 5f) for ss_-27_-3_ds_-2_+143_-R.

#### Image processing of ss_-27_-8_ds_-7_+143_-R, ss_-27_-13_ds_-12_+143_-R, dsR_-67_+113_(poly-dT15_-17_-3_)

It was evident from the processing of earlier datasets that the docking of TE domain indicating localized melting of *oriT* could be visualized from high resolution 2D classes and was observable even in initial 3D maps obtained from *ab initio* models in cryoSPARC. Therefore, for the next constructs tested for *oriT* melting, the data processing was carried out only till we obtained initial 3D maps. Briefly, pre-processing of the datasets were carried out as above, followed by 2D classification and a round of *ab initio* classification with refinement of the best class yielding low resolution 3D maps. For comparison, initial 3D maps obtained during early processing steps of ds_- 27_+143_-R and ss_-27_+8_ds_+9_+143_-R at a similar resolution are shown (Fig. 3d,e). The data collection statistics, resolution and initial and final particles obtained during processing are given in ED Table 2.

All maps used for model building were sharpened using DeepEMhancer^57^ and local resolution estimation was performed using cryoSPARC.

### Model building and refinement

Atomic model building into the 3.78 Å ds_-27_+143_-R locally refined map was initiated by fitting the crystal structure of the IHF-DNA module (PDB: 1IHF) using UCSF Chimera^58^. Regular B-form DNA generated by Web 3DNA^59^ was used as an initial model for the DNA arms and modified to fit the trajectory and helical pitch of the visible DNA density, using all-atom refine in COOT v0.9.8^60^ with ProSMART restraints^61^. Three copies of a model of TraY generated using AlphaFold2 (AF) ^62^ were fitted into the three TraY densities visible in the cryoEM map using UCSF Chimera. AlphaFold2 predicted model of F TraI was similar to the experimental structure of its homologue R1 TraI in its helicase mode. From the F TraI AF model, the VH_2A+2B/2Blike_ sub-domain structure was used for fitting into the corresponding density in the map. The complete model was then subjected to a few rounds of iterative manual rebuilding in COOT and refinement in PHENIX^63^.

The low resolution of the map compounded by motion of the DNA arms as predicted from the 3D variability analysis in cryoSPARC, led to some ambiguity in the DNA base density. Although individual bases could be resolved, confident assignment and differentiating between purines and pyrimidines in the *oriT* density remained challenging. To assign the DNA sequence, we decided to take advantage of the three known TraY binding sub-sites described by the Schildbach group (16). Therefore, starting from the three known TraY sites we manually updated the DNA sequence in COOT following the *oriT* DNA register. This assignment was only approximate, and the sequence was later reassigned guided by the ss/ds heteroduplex relaxosome maps which were of much higher quality in the DNA region (see below).

Further fitting of individual residues into the density were carried out by a combination of manual building using COOT and MDFF using ISOLDE^64^. The model was inspected for correct rotamer/Ramachandran outliers and clashes in ISOLDE. Real-space refinement of the atomic model of ds_-27_+143_-R was carried out in PHENIX with Ramachandran, secondary structure and nucleic-acid restraints applied.

To build the ss_-27_+8_ds_+9_+143_-R model into the corresponding 3.45 Å locally-refined map, the *de novo* built ds_-27_+14_-R model was initially fitted into the density using UCSF Chimera. The crystal structure of the F TraI trans-esterase (TE) domain bound to its cognate ssDNA (PDB ID: 2A0I) was used as a model for fitting in the ordered TE density of the map. The higher resolution of the map and the stability of DNA arms resulted in an improved DNA density where the purines and pyrimidines were now clearly resolved. Therefore, the DNA sequence was accurately reassigned in the model manually using COOT. Individual protein and DNA residues were rebuilt into densities by iterative rounds of a combination of manual building in COOT and MDDF in ISOLDE. The model was inspected to correct rotamer/Ramachandran outliers and clashes using ISOLDE followed by refinement in PHENIX with restraints applied.

The completed atomic model for ss_-27_+8_ds_+9_+143_R without the TE-ssDNA was then used to further improve the ds_-27_+14_-R model. After fitting into the ds_-27_+14_-R map, it was rebuilt using COOT and ISOLDE with iterative rounds of real space refinement with restraints applied in PHENIX, to yield the final model for ds_-27_+143_-R.

The ss_-27_+8_ds_+9_+143_-R model was also used as an initial model for the ss_- 27_+8_ds_+9_+143_RΔ_AH+CTD_, ss_-27_+8_ds_+9_+143_RΔ_TraM_, ss_-27_+3_ds_+4_+143_R and ss_-27_-3_ds_-2_+143_R, rebuilt using COOT and ISOLDE with iterative rounds of real space refinement with restraints applied in PHENIX, to yield the final models for each of the complexes. These atomic models were nearly identical to ss_-27_+8_ds_+9_+143_-R with an average rms deviation of 0.738 Å for the Cα superposition.

For all the models, regions of poor Cα backbone density were subsequently deleted, and the side chains removed for areas with poor densities for side chains. The models were validated using Molprobity^65^. The statistics for model building, refinement and validation are reported in ED Table 1b.

For fitting models of VH_NTD_, VH_1A_, AH and TraM to generate the ss_-27_+8_ds_+9_+143_- R global model, the unsharpened global ss_-27_+8_ds_+9_+143_-R 3.77 Å map was used. Initially, the refined ss_-27_+8_ds_+9_+143_-R structure was fitted in the global map. To generate a model for F TraM bound to DNA, an AF model of F TraM was first superposed onto the pED208 TraM:*sbmA* structure (PDB ID: 3ON0) with the CTDs as a guide. The NTDs of F TraM were adjusted to position on the NTDs of the DNA bound conformation of pED208 TraM to generate a model of F TraM bound to pED208 *sbmA*. This F TraM:pED208 *sbmA* model was docked as a rigid body into the map using UCSF chimera Fit in Map tool (cross-correlation 0.80 of map to model). The ends of the ss_-27_+8_ds_+9_+143_-R DNA proximal to TraM was extended by two base pairs, joined to pED208 *sbmA* DNA and the sequence was updated in the F TraM:pED208 *sbmA* model to match the *oriT* sequence by mutating DNA residues in COOT. From the AF generated model of F TraI, VH_NTD_ (residues 307-385) and VH_1A_ (residues 383-564) subdomains were separated and individually fitted in their densities in the map (correlation coefficients of 0.87 for VH_NTD_ and 0.89 for VH_1A_). Next, Isolde was used to flexibly refine these models with adaptive distance restraints implemented. The models were then merged with the VH_2A+2B/B-like_ sub-domain to yield the complete VH domain.

For the AH domain, sub-domains of the F TraI AF model were fitted individually. The fitting was initiated with AH_1A_ for which the density exhibited partial secondary structure features and the AH_1A_ model (residues 934-1126) could be fitted with a correlation of 0.87. AH_NTD_ (residues 835-933) was fitted in its density (correlation coefficient of 0.91) and both the models were flexibly refined in Isolde with adaptive distance restraints. Next the CTD (residues 1460-1630) model was rigid body fitted in the map with a correlation of 0.84, and flexible fitting was carried out in Isolde with restraints applied. The AH_2A+2B/B-like_ model (residues 1127-1459) was manually fitted in the density as this region was fragmented and poorly defined. The fitting was done such that the N-terminus and C-terminus of the AH_2A+2B/B-like_ model was proximal to the C terminus of AH_1A_ model and the N terminus of CTD, respectively. AH_NTD_, AH_1A_ and AH_2A+2B/B-like_ models were joined to give the full AH domain. This gave us the composite global model of a fully assembled relaxosome. The models thus assembled gave rise to minimal clashes in the structure and the fitting of sub-domains as described above resulted in the corresponding N and C termini being in proximity such that they could all be merged confidently. Note that the side chains for VH_NTD_, VH_1A_, AH and CTD were removed in the final model.

### Interactions, analysis, visualization and representation

Interactions were calculated using the PDBePisa server^66^. Domain movements were analysed using the DynDom program^67^. CryoEM maps and models were visualized and figures prepared using UCSF Chimera^58^, UCSF ChimeraX^68^ and PyMOL^69^.

### Construction of p99I^+^ mutant derivatives

Double- or triple- site mutations in F *traI* were generated by cutting p99I^+^ with Eco0901 to remove a 1.4 kb fragment of TraI or in case of variant S73E W659 with KpnI/Eco0901 to remove 3 kb. Forward primer MutI fw and the reverse primer MutI rev were used to create all mutant alleles except mutant S73E W659E which required forward primer MutI17fw (ED Table 4c). Mutant-specific primer pairs (ED Table 4d) containing the mutations in their overlap region were used together with these forward/reverse pairs to generate fragments suitable for error free joining via seamless assembly of multiple DNA Fragments [NEBuilder HiFi DNA Assembly Kit (NEB)]. *E. coli* DH5α cells were transformed with the ligation products. Oligonucleotide synthesis and Sanger sequencing of the mutated variants was performed by Microsynth Austria. **Bacterial conjugation**

Conjugative plasmid transfer was performed as described in^70^. The conjugation deficient phenotype of *E. coli* MS411 [pOX38Δ*traI*] donor cells was complemented in trans with wild-type [p99I^+^] or mutant *traI* alleles. Overnight cultures of donor strains were diluted to an optical density at 600 nm (OD_600_) of 0.005 in 900 µl antibiotic-free LB medium and incubated for 1 h at 37°C before 100 µl of recipient culture (OD_600_ of 2) was added. Conjugation was allowed to proceed for 1 h at 37°C and then stopped by vortexing for 1 min. Transconjugants were selected on plates containing 25 µg/ml Sm and 40 µg/ml Km. Conjugation frequencies were expressed as transconjugants per donor cell (ED Table 3).

### Statistical analysis

Statistical analysis was performed using GraphPad prism version 10. Comparisons of conjugation frequencies of wild-type TraI and mutated variants for each interface hub was performed by ordinary one-way ANOVA with Holm-Šídák’s multiple comparisons test; P<0.05 was considered significant (ED Table 3).

## Extended Data Tables and Figures

**ED Fig. 1.**
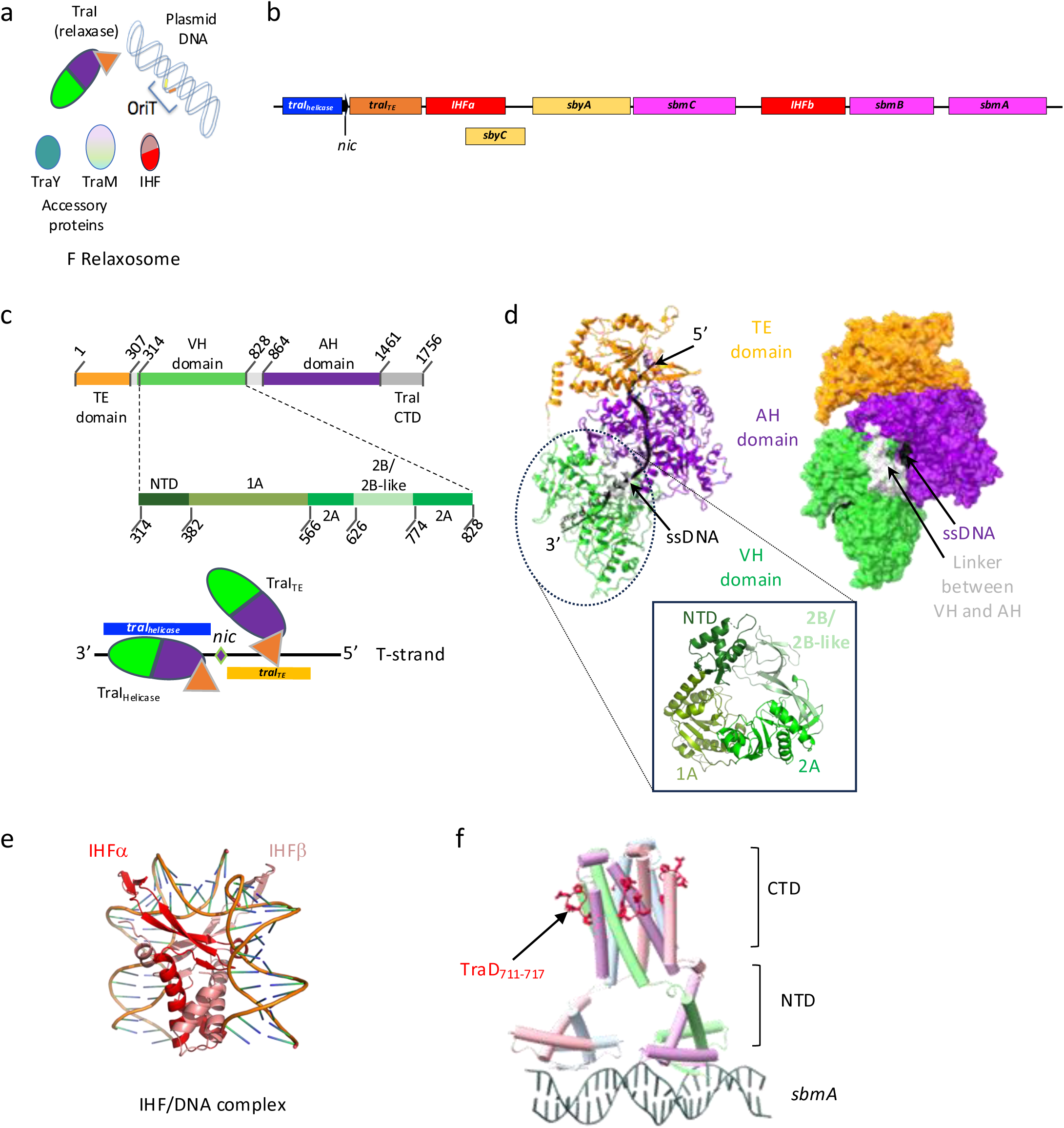
Prior knowledge on the structure of relaxosome components. a, F plasmid relaxosome components. b, *oriT* region of the F plasmid. The binding sites for each relaxosome protein as defined by Ilangovan *et al*., Frost *et al*. and Lum *et al*.^6-8^ for IHF (*IHFa* and *IHFb*), TraM (*sbmA-C*), TraY (*sbyA* and *sbyC*), and TraI (*traI_helicase_* and *traI_TE_*) are indicated under *oriT_316_* using boxes colour-coded red for IHF, yellow for TraY, dark blue, and orange for TraI_helicase_ and TraI_TE_, magenta for TraM, and black for the *nic* site. c, Domain structure of TraI (top and middle) and the two binding modes of TraI. Top: TraI domains mapped on primary sequence. TE, VH, AH, and CTD are shown in orange, green, violet, and grey, respectively. Boundary residue numbers are shown. Middle: sub-domains of VH mapped on primary sequence. Boundary residues for NTD, 1A, 2A, and 2B/2B-like are indicated. Bottom: TraI binds sequence of the T-strand 3’ to *nic* in its compact, protease-resistant, helicase mode (termed “TraI_helicase_”), while it binds sequences of the T-strand 5’ to *nic* in its TE mode (“TraI_TE_”). d, Structure of TraI_helicase_ (PDB ID 5N8O). This structure was obtained using a 22-mer ssDNA (in magenta). Upper left and right: two views of TraI, one in ribbon and one in surface representations. Various domains and the DNA are labelled. Linker between VH and AH is shown in grey. The CTD was not part of this structure. Bottom: structure of the VH domain showing the location of the various subdomains. Color-coding is as in panel c. e, Structure of *Escherichia coli* IHF bound to an IHF-binding site on phage lambda DNA (PDB ID 1IHF). IHF binding generates a bend of > 160° on dsDNA. f, Composite model of pED208 TraM bound to *sbmA* (PDB ID 3ON0) and F TraD_711-717_ (PDB ID 3D8A). Protein and DNA are shown in ribbon, and TraD_711-717_ residues in red sticks. Each chain in the TraM tetramer is shown in a different colour.

**ED Fig. 2.**
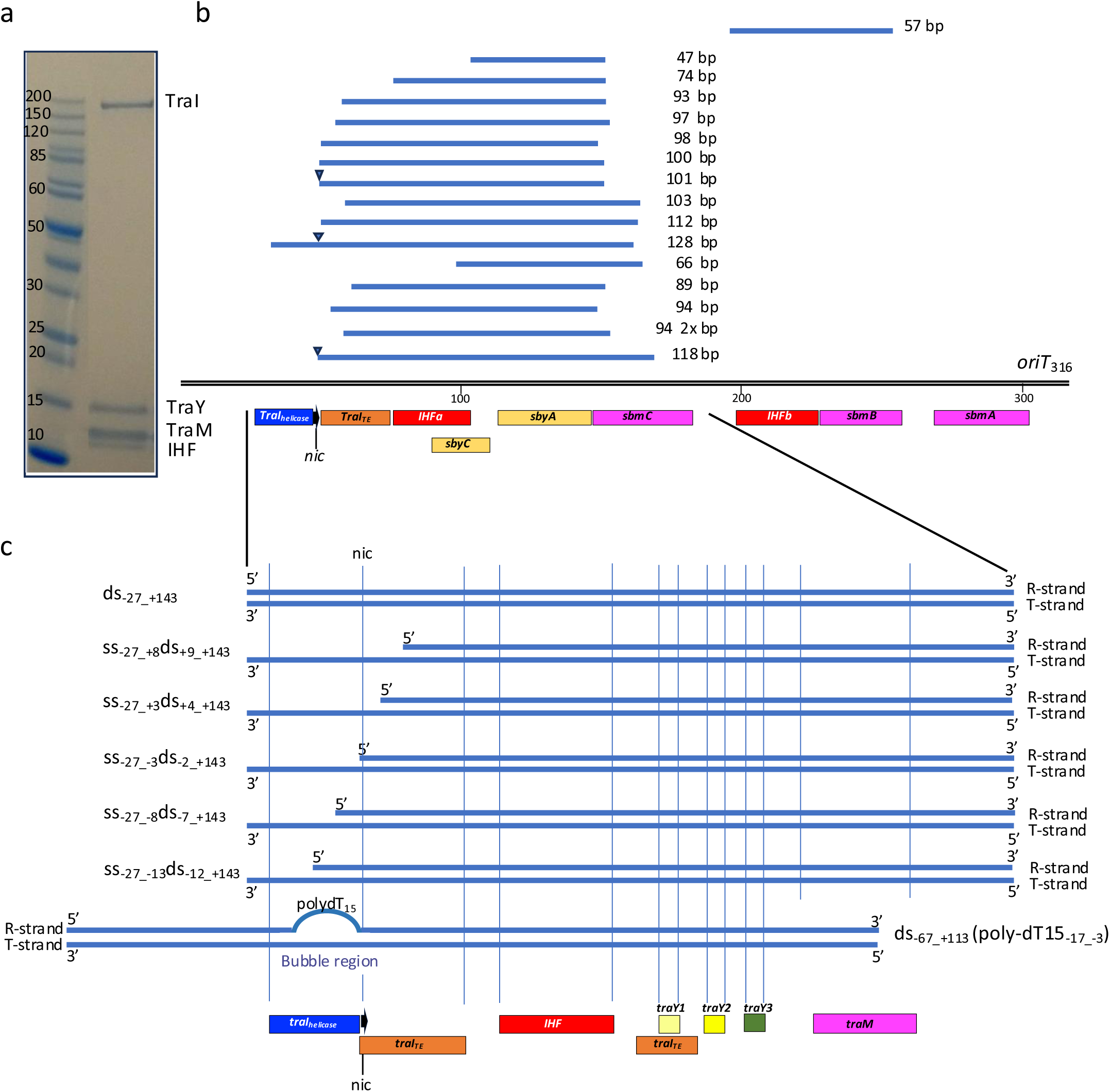
Biochemical characterisation of the F plasmid relaxosome a, SDS-PAGE gel of the relaxosome. Purified TraI, TraY, IHF, and TraM were mixed with *oriT_316_* and unbound components were excluded by size-exclusion chromatography, leading to a purified relaxosome. Molecular Weight markers are labelled. b, *oriT* sequences protected from DNase I nuclease cleavage. The sequenced fragments represented by blue lines are mapped against *oriT_316_*. The length of each fragment is shown. The various binding sites for each relaxosome protein as defined by Ilangovan et al. (2017) for TraI^6^ and by Frost *et al*., and Lum *et al*.,^7,8^ for IHF, TraM, and TraY are shown under *oriT_316_* using boxes colour-coded red for IHF, yellow for TraY, dark blue and orange for TraI_helicase_ and TraI_TE_, and magenta for TraM. When present in the DNA fragment, the position of the *nic* site is shown by a black filled triangle. c, DNAs used in this study. The DNAs are labelled according to the following notation: DNAs are numbered relative to the *nic* site on the T-strand. Therefore, *nic* serves as the origin, with the bases or base pairs 3’ to it being negatively numbered, and the bases and base pairs 5’ to it being positively numbered. This notation provides instant recognition of the position of *nic* in the DNA used. Furthermore, “ss” and “ds” DNA is used to indicate which region of *oriT* is single-stranded or double-stranded in the DNA used. For example, ss_-27_+8_ds_+9_+143_ is a DNA which is single-stranded from base +8 to -27 and double-stranded from +9 to +143. The various binding sites for each relaxosome proteins as discovered in this study are shown underneath. Box colour-coding is as in panel b except for the three TraY molecule train which are coloured pale yellow, bright yellow, and olive green for TraY1, TraY2, and TraY3, respectively. The position of the *nic* site (black arrow) is shown.

**ED Fig. 3.**
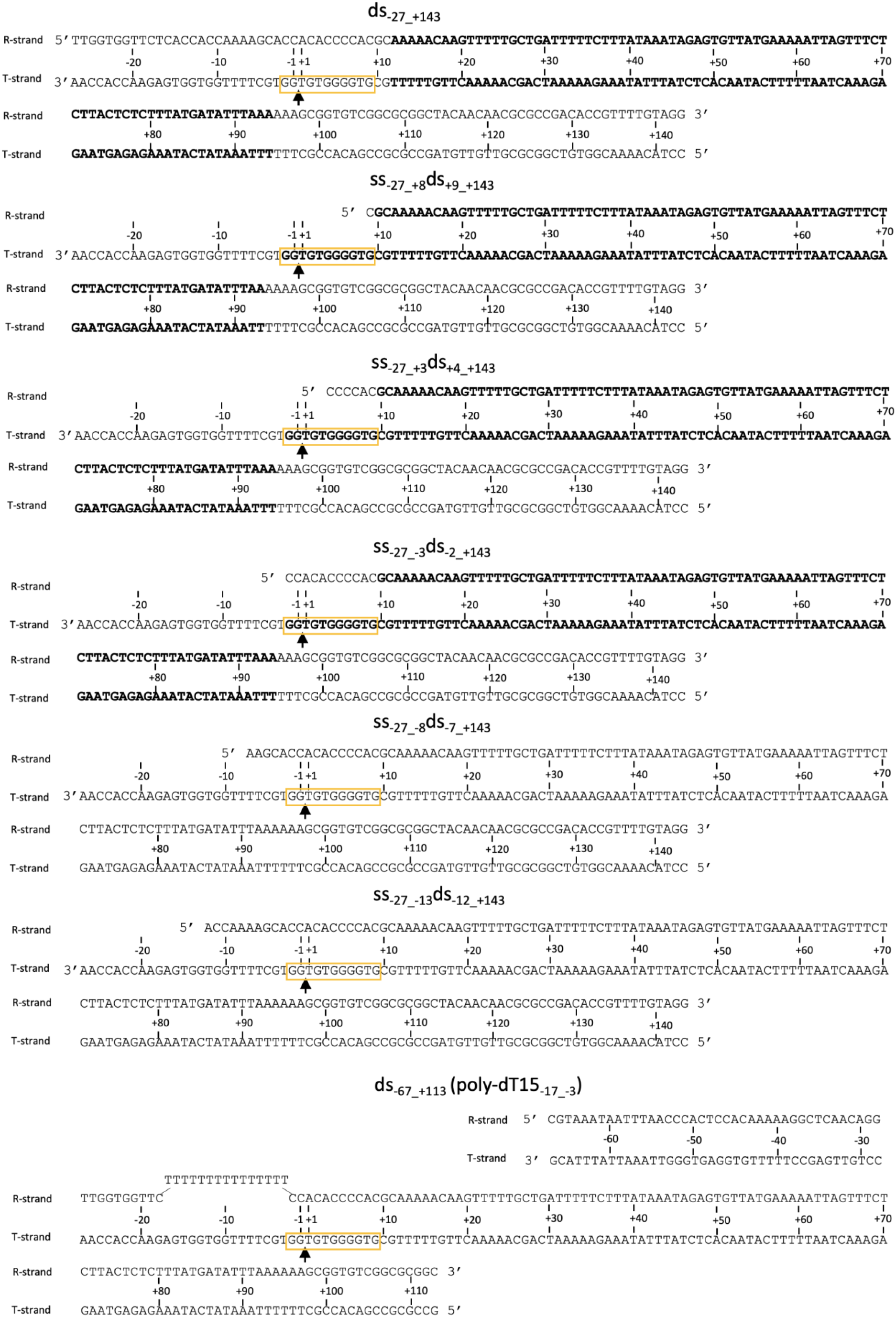
DNAs used in this study. Naming and numbering is as in ED Fig. 2c. *nic* is indicated by an arrow. The region of DNA for which a model could be derived from the density is shown in bold. Orange box locates the TE-binding site on the T-strand between bp_-2_ and bp_+9_.

**ED Fig. 4.**
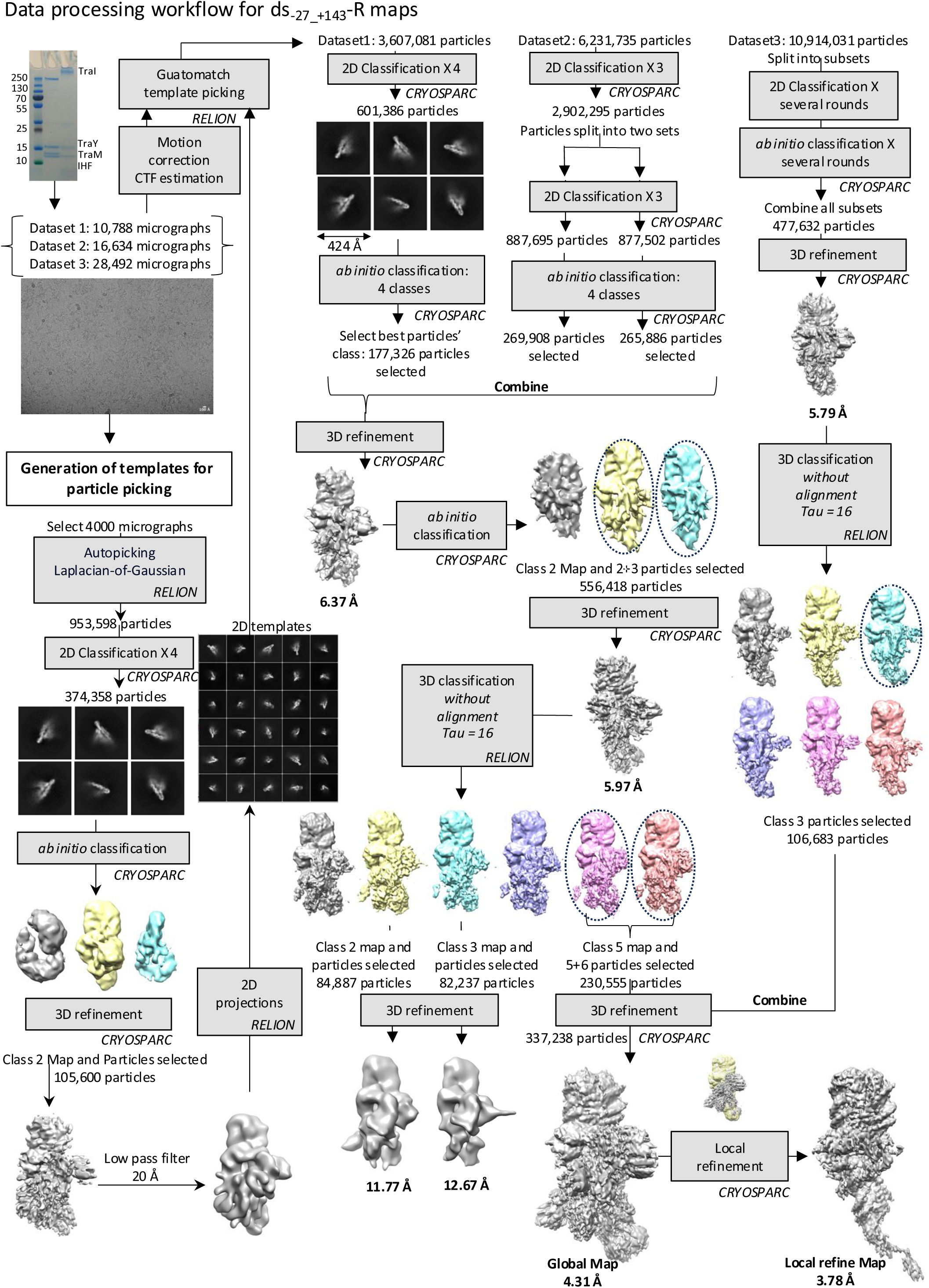
Image processing workflow for the cryo-EM structure determination of the ds_-27_+143_-R complex. This being the first to be determined for any relaxosome structure, the *ab-initio* determination workflow is provided.

**ED Fig. 5.**
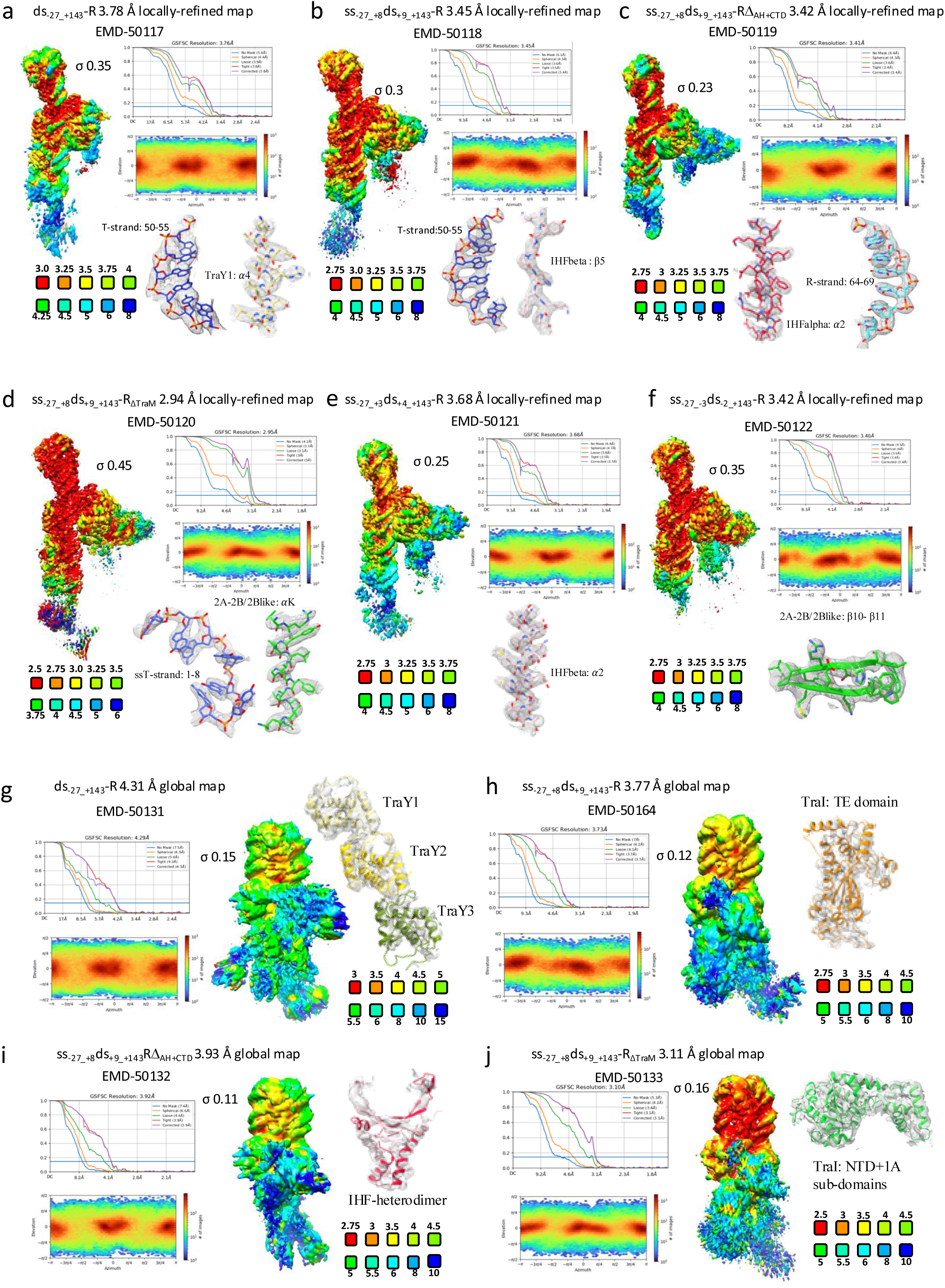
Cryo-EM maps used in this study. Maps are termed “locally-refined” or “global”, depending on whether local refinement using a mask encompassing the DNA, the three TraYs, IHF, TraI VH_2A+2B/2B-like_ and TraI_TE_ has been applied or not, respectively. For each map, the density coloured by local resolution, the average resolution derived from Fourier Shell Correlation (FSC), the angular distribution and a representative region of the electron density map with the final model of the relaxosome built in it (in stick representation colour-coded as in Fig. 1e) are shown. Local resolution was calculated using CRYOSPARC (FSC cut-off 0.143) and coloured as indicated in the scale below the map. For each map, FSC plots show curves for correlation between 2 independently refined half-maps with no mask (blue), spherical mask (orange), loose mask (green), tight mask (red) and corrected (purple). Cut-off 0.143 (blue line) was used for resolution estimation. a, ds_-27_+143_-R locally-refined 3.78 Å map; b, ss_-27_+8_ds_+9_+143_-R 3.45 Å locally-refined map; c, ss_-27_+8_ds_+9_+143_-RΔ_AH+CTD_ 3.42 Å locally-refined map; d, ss_-27_+8_ds_+9_+143_-RΔ_TraM_ 2.94 Å locally-refined map; e, ss_-27_+3_ds_+4_+143_-R 3.68 Å locally-refined map; f, ss_-27_-3_ds_-2_+143_-R 3.42 Å locally-refined map; g, ds_-27_+143_-R global 4.31 Å map; h, ss_-27_+8_ds_+9_+143_-R 3.77 Å global map; i, ss_-27_+8_ds_+9_+143_-RΔ_AH+CTD_ 3.93 Å global map; j, ss_-27_+8_ds_+9_+143_-RΔ_TraM_ 3.11 Å global map.

**ED Fig. 6.**
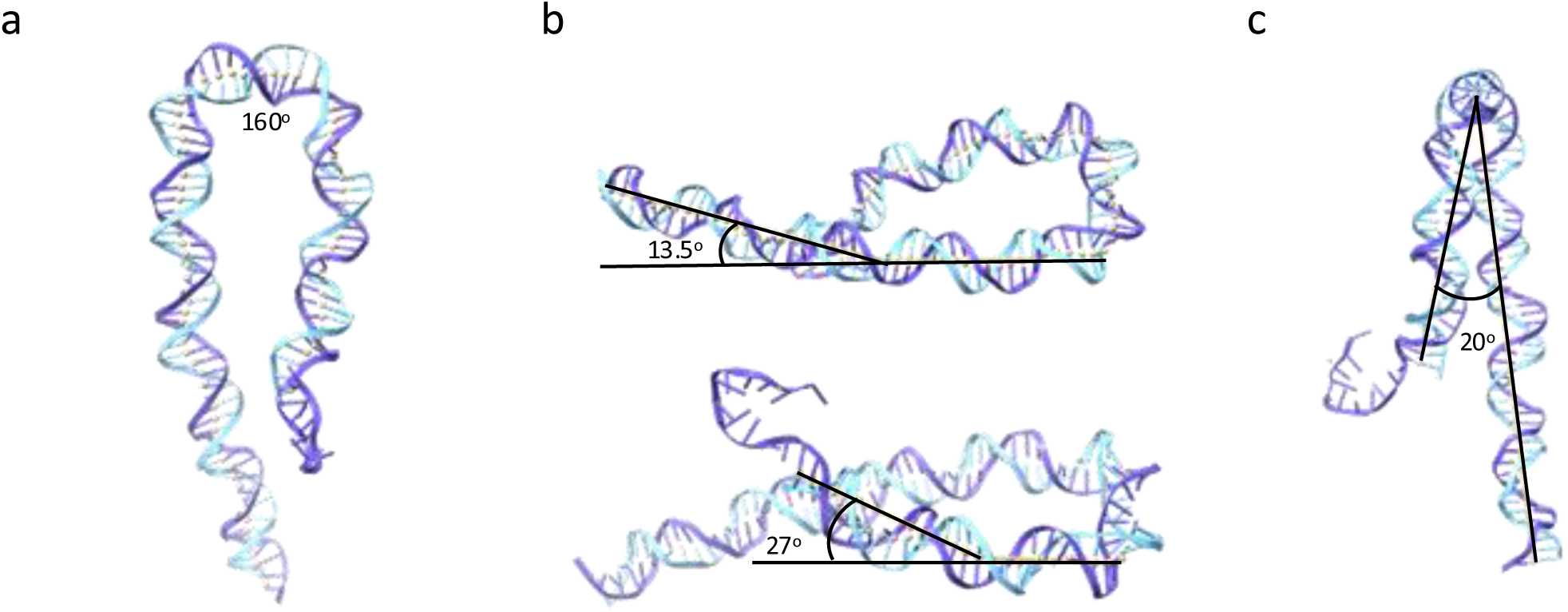
Assessment of dsDNA bending from ss_-27_+8_ds_+9_+143_-R structure. a, Bending induced by IHF binding evaluated as in Rice *et al.*^9^. b, Bending within the two DNA hairpin arms. Top panel: Bending triggered by the binding of the TraY train on the TraY arm of the DNA hairpin. Bottom panel: Bending in the *nic* arm is the result of sequence-induced bending between base pairs +20 and +34^10^ and binding of VH_2A+2B/2B-like_. Angles are reported. c, Bending between the two DNA hairpin arms. When the DNA is viewed such that the IHF bend is perpendicular to the plane of view, the arms of the hairpin make an angle of 20°, seemingly moving away from each other.

**ED Fig. 7.**
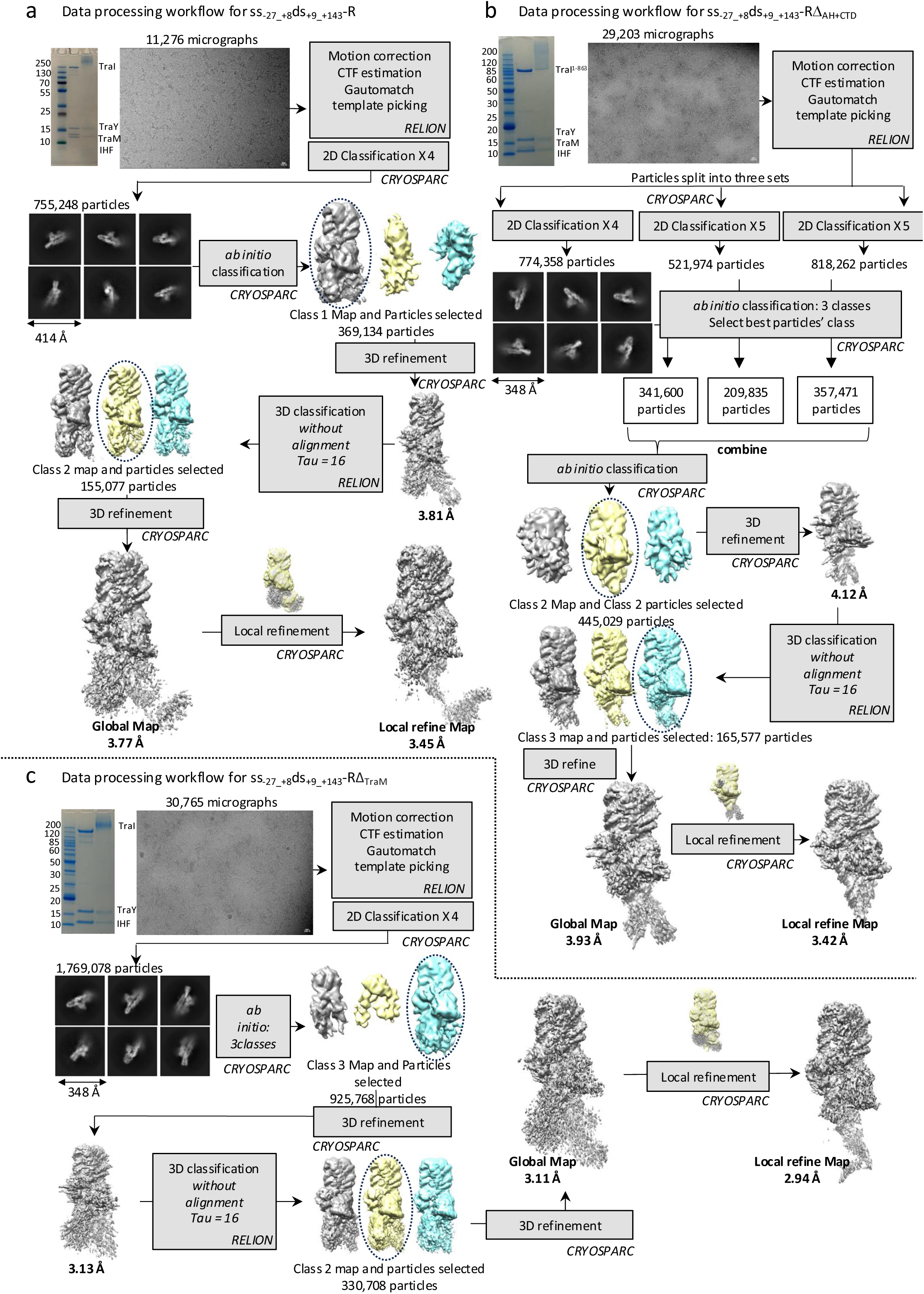
Image processing workflow for the cryo-EM structure determination of the (a) ss_-27_+8_ds_+9_+143_-R, (b) ss_-27_+8_ds_+9_+143_-RΔ_AH+CTD_ and (c) ss_-27_+8_ds_+9_+143_-RΔ_TraM_. Workflow for the maps shown in ED Fig. 5e (ss_-27-+3_ds_+4_+143_-R 3.68 Å locally-refined map) and 5f (ss_-27_-3_ds_-2_+143_-R 3.42 Å locally-refined map) are not provided as they are essentially similar to that provided for ss_-27_+8_ds_+9_+143_-R.

**ED Fig. 8.**
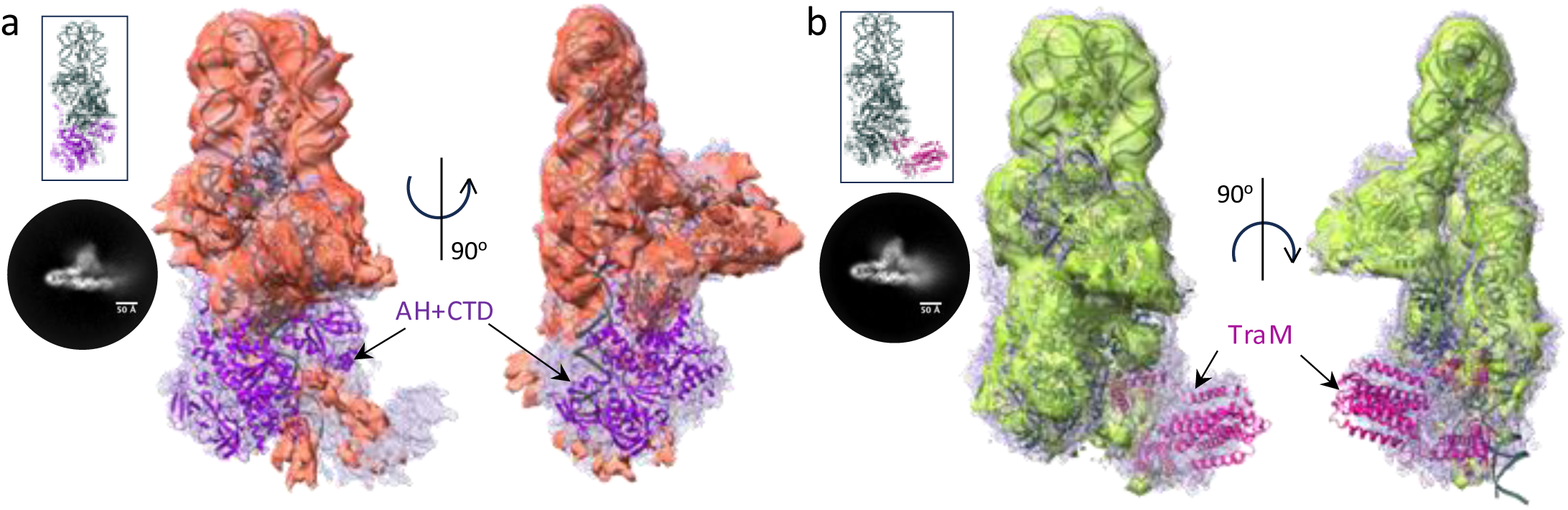
Use of maps where parts of the relaxosome have been deleted to generate the complete structure of the relaxosome. a, Superposition of the unsharpened global maps of ss_-27_+8_ds_+9_+143_-R (in light blue mesh) and ss_-27_+8_ds_+9_+143_-RΔ_AH+CTD_ (in light red surface). Maps details are reported in ED Figs. 5h and 5i, respectively. A representative 2D class is shown on the left. The structure of ss_-27_+8_ds_+9_+143_-R is in dark grey ribbon. A model of TraI AH and CTD (in violet ribbon) is shown fitted in the additional density observed in ss_+8_-27_ds_+9_+143_-R, but not in ss_+8_-27_ds_+9_+143_-RΔ_AH+CTD_. Two orientations, 90° apart, are shown at left (same orientation in inset) and right. The lack of resolution for TraI AH and CTD is not surprising. Firstly, TraI is an extremely flexible protein. It unfolds readily to pass through the pilus during conjugative transfer and it is known to be unusually susceptible to mild-proteolytic cleavage either in its unbound form or bound to its TE ssDNA-binding site (*traI_TE_* in ED Fig. 2b)^6,11^. It is known to be ordered only in its helicase-binding mode^6,12^. Moreover, both TraI AH and CTD together with TraM are located in a region of the DNA that is itself ill-defined, indicating flexibility of the DNA in this region. b, Superposition of the unsharpened global maps of ss_-27_+8_ds_+9_+143_-R (in light blue mesh) and ss_-27_+8_ds_+9_+143_-RΔ_TraM_ (in light green surface). Map details are reported in ED Figs. 5h and 5j, respectively. A representative 2D class is shown on the left. The structure of ss_-27_+8_ds_+9_+143_-R is in dark grey ribbon. A model of TraM (in magenta ribbon) is shown docked in the additional density observed in ss_+8_-27_ds_+9_+143_-R but not in ss_+8_-27_ds_+9_+143_-RΔ_TraM_. Two orientations, 90° apart, are shown at left (same orientation in inset) and right. To validate the TraM location, we also generated an extended model of the DNA that includes *sbmC* and observed that *sbmC* locates in the density region attributed to TraM. Moreover, the molecular interactions of TraM within this region of DNA are very similar to those identified previously by hydroxyl radical foot printing thereby providing another layer of confidence that the density for TraM has been properly ascribed (Fig. 2g and ED Fig. 10c)^13^.

**ED Fig. 9.**
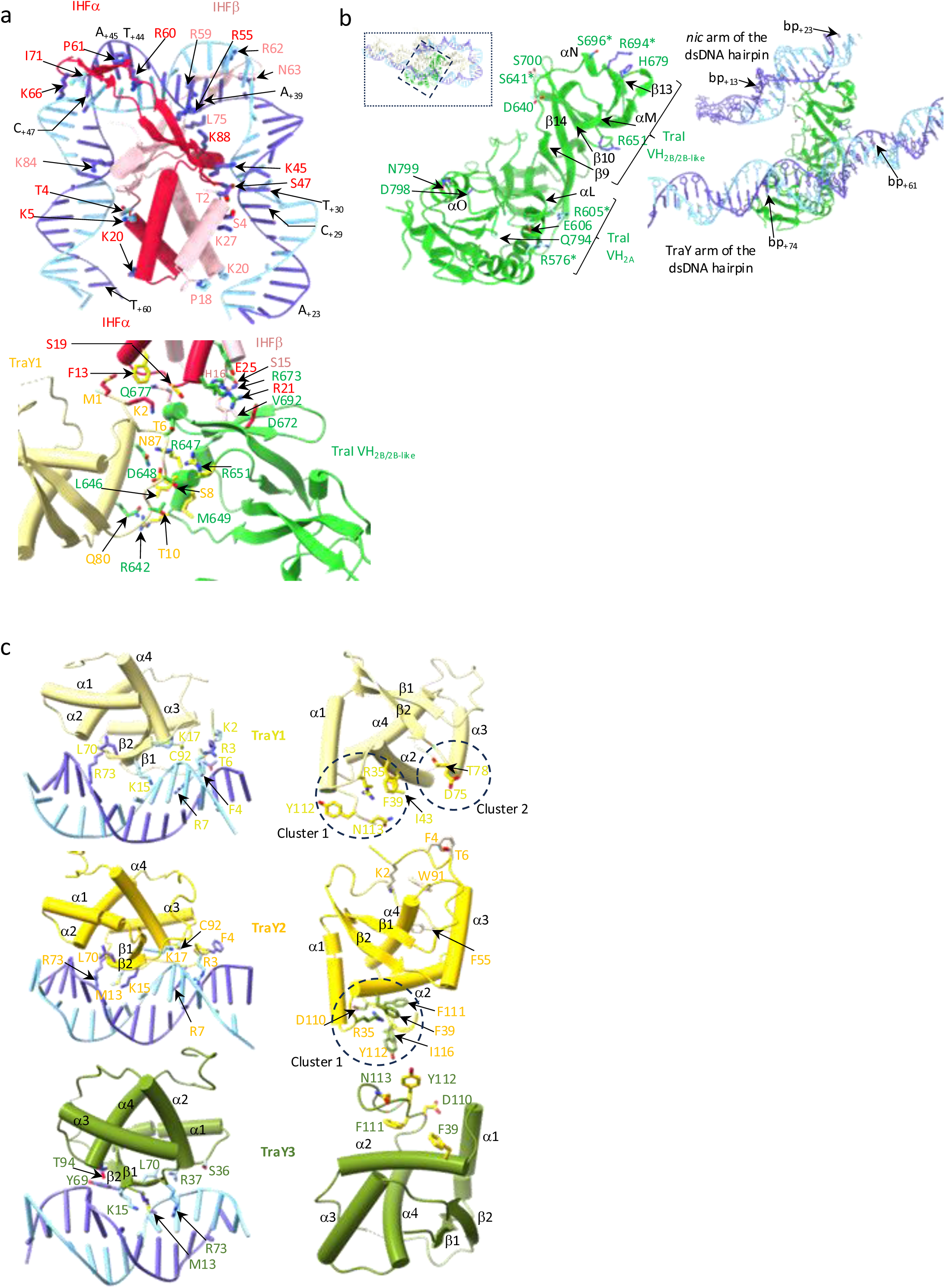
Details of the fully-assembled ss_-27_+8_ds_+9_+143_-R structure. a, Details of interactions in Hub1. Representation, colour-coding, and labelling is as in Fig. 2b, except for interacting residues which are shown in stick representation and colour-coding according to the colour of the DNA or protein they interact with. b, Details of interactions between TraI VH_2A+2B/2B-like_ and DNA. Left: TraI VH_2A+2B/2B-like_ residues involved in dsDNA-binding are shown in stick representation colour-coded in light and dark blue when interacting with the R- or T-strand, respectively. Right: same view as at left, with DNA and binding-site boundaries shown. The two arms of the dsDNA hairpin are indicated. c, Details of interactions in Hub2. Representation, colour-coding, and labelling is as in Fig. 2d, except for interacting residues which are shown in stick representation and colour-coded according to the colour of the DNA strand (left panel) or protein (right panel) they interact with. Left: TraY-DNA interactions for TraY1 (top), TraY2 (middle), TraY3 (bottom). Labelling of residues is as for the protein but their stick colour is that of interacting DNA strand, light blue for the R-strand and dark blue for the T-strand. Right: protein-protein interactions for TraY1 (top), TraY2 (middle), and TraY3 (bottom). Labelling of residues are as at left but stick colour is that of the protein the residue interacts with. TraY’s binding to DNA is typical of RHH dimers with the β1 and β2 strands interacting with the major groove of the DNA binding site^14^. But binding of TraY1 and TraY2 extends to the minor groove as well with residues at the extended N-terminus (before β1) contributing to minor groove binding. The third TraY (TraY3) binding site on DNA was predicted to be inverted and indeed TraY3 binds in an inverted orientation compared to TraY1 and TraY2. This agreed with a study by Lum *et al*.^8^ which showed TraY binding to three sub-sites on *sbyA* (ED Figs. 1b and 2c). As a result, the N-terminal residues involved in minor groove binding in TraY1 and TraY2 are no longer involved. Also, residues in α1 and α2, which in TraY1 and TraY2 are located away from the DNA binding site, are now involved in DNA-binding. TraY1 interaction with TraY2 involves two clusters of residues: the first at the C-terminus and in α1, α2, and α1-α2 (circle labelled “cluster 1”) and another in β2-α3 and α3 (circle labelled “cluster 2”). Both regions contact TraY2 via the N-terminus of TraY2 but also its α2-β2 loop and α3. However, TraY2 interacts completely differently with TraY3 because TraY3 is inverted due to binding to the inverted DNA repeat. In fact, the TraY2-TraY3 interaction involves the same parts of the two proteins, namely cluster 1 mentioned above. Thus, cluster 1 of TraY2 and TraY3 face head to head in a manner that is conducive to complementary interactions between hydrophobic residues (F111, F39 for example) and charged residues to make ion pairs (D110, R35 for example).

**ED Fig. 10.**
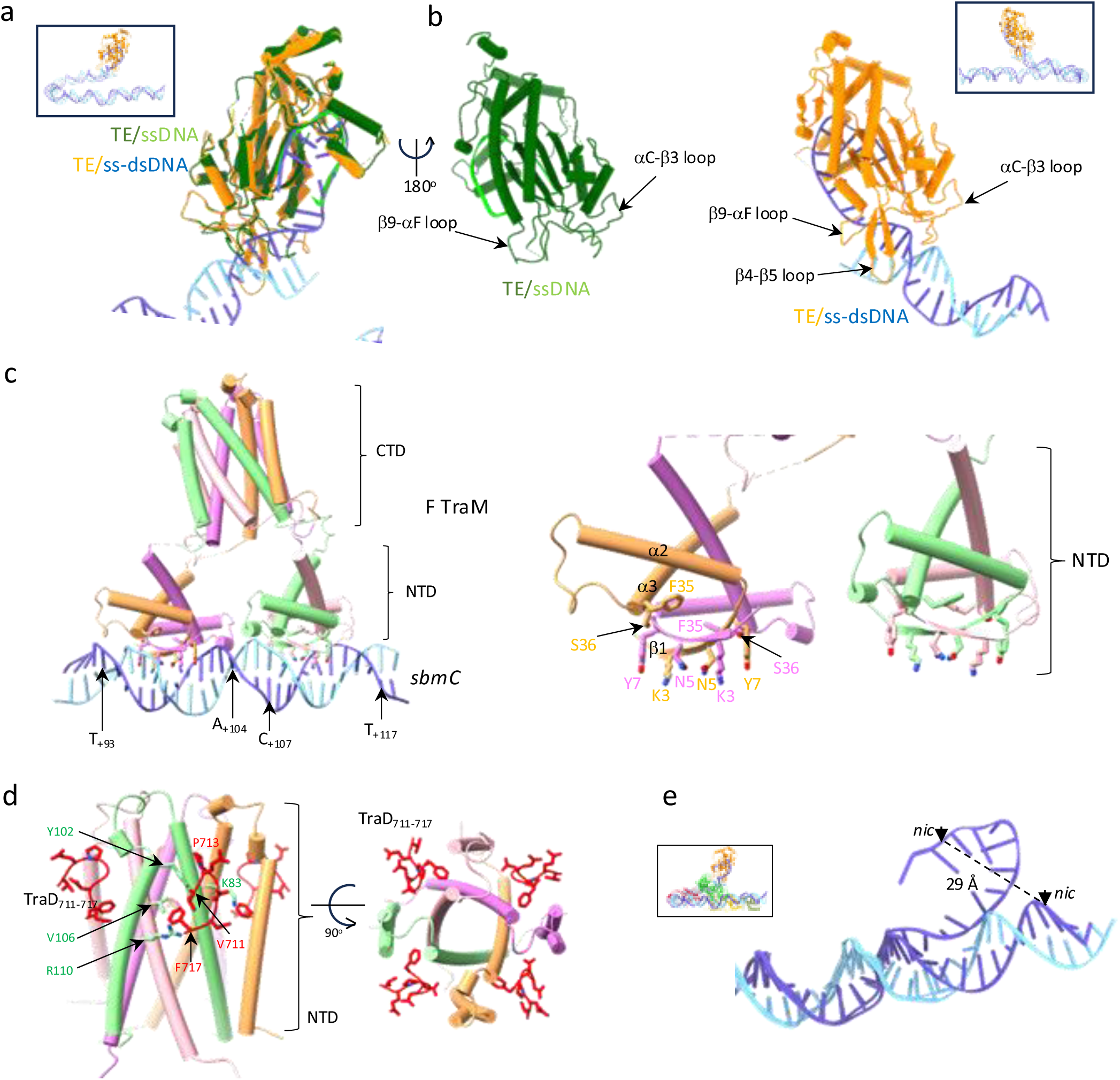
Interactions of TraI, TraM with TraD and *oriT* conformational change. a, Superposition and comparison of TraI TE bound to ss_-27_+8_ds_+9_+143_ DNA (TE/ss-dsDNA) and the previous crystal structure of TraI TE bound to ss_+9_-2_ determined previously (TE/ssDNA; PDB ID 2A0I). RMSD in Cα position of 0.649 Å. b, Secondary structures being re-organised or ordered upon binding of the ds region of ss_-27_+8_ds_+9_+143_ in the ss_-27_+8_ds_+9_+143_-R relaxosome structure. Left: TE/ssDNA structure. Right: TE/ss-dsDNA structure. The part of the structure undergoing conformational changes are labelled. c, TraM binding to *sbmC*. Protein and DNA are shown in ribbon, colour-coded in four different colours for the 4 different TraM chains. R- and T-strand are in light and dark blue ribbon, respectively. Left: overall view of TraM binding. Boundary base pairs for the two TraM NTD dimer binding sites are indicated. Right: Residues involved in binding. Only residues for the binding site at left are labelled. d, TraD_711-717_ binding to TraM. The co-crystal structure of TraM-TraD (PDB ID: 3D8A) was used with the same colour-coding for TraM as above. TraM is shown in ribbon except for the residues involved in binding which are shown in stick. TraD_711-717_ is shown in stick colour-coded red. Residues involved in interactions are labelled. e, The ss_-27_+8_ds_+9_+143_ DNA in the ss_-27_+8_ds_+9_+143_-R structure superposed on the ds_-27_+143_ DNA in ds_-27_+143_ -R structure where the ds_-27_+143_ DNA has been extended to include the *nic* site. This superposition highlights the very large motion undergone by *nic* and its DNA region upon binding TraI TE.

**ED Table 1a.**
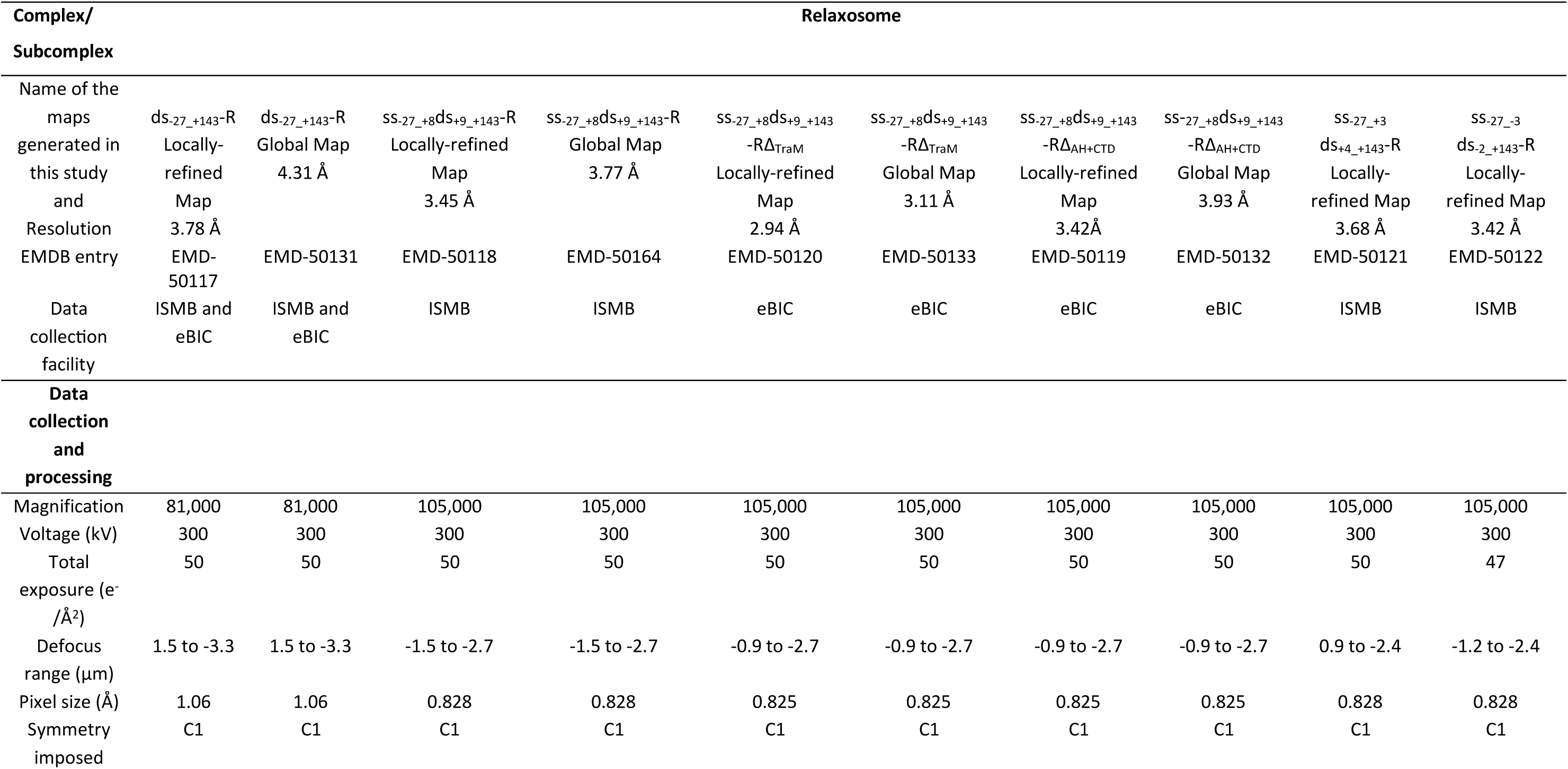
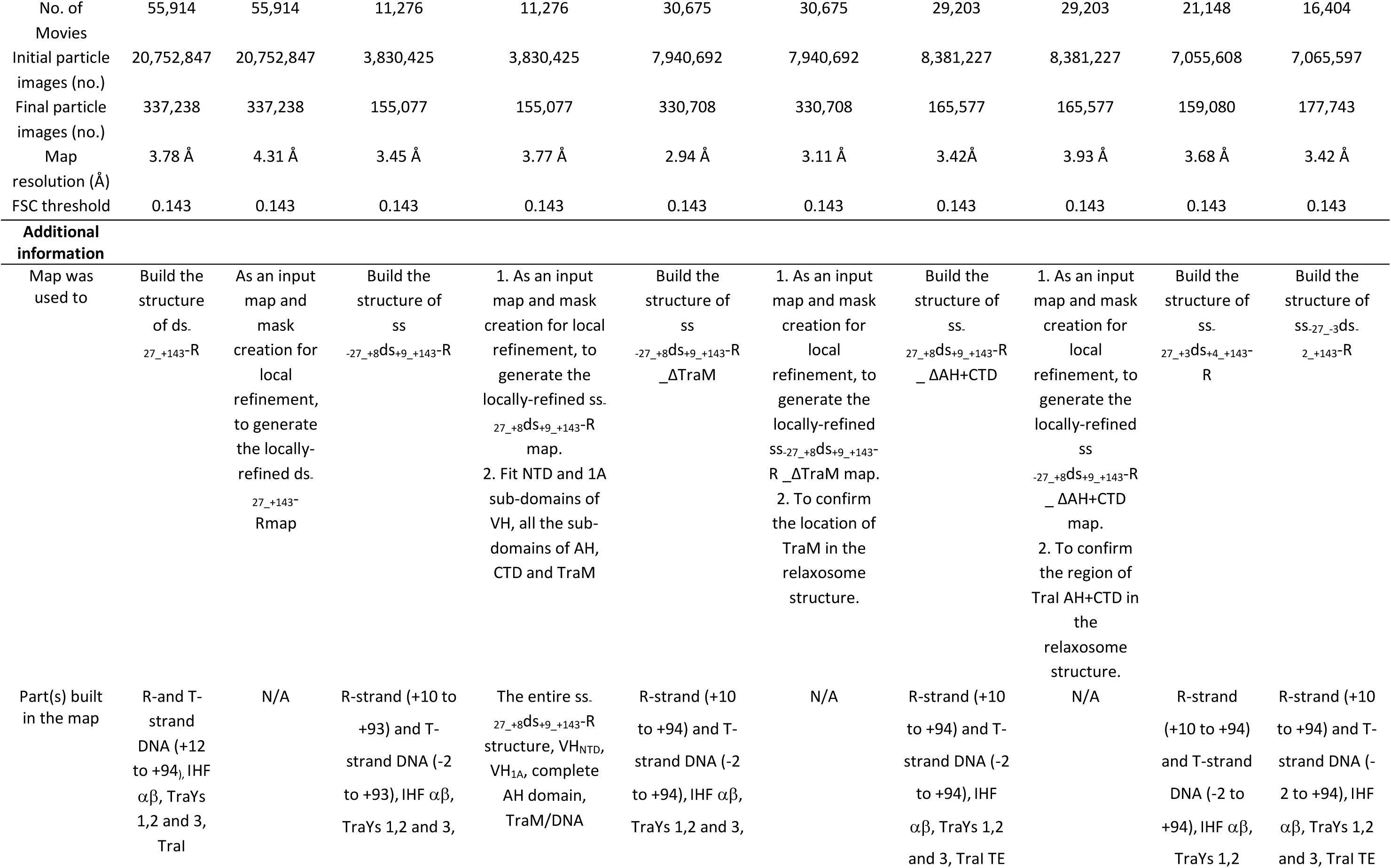
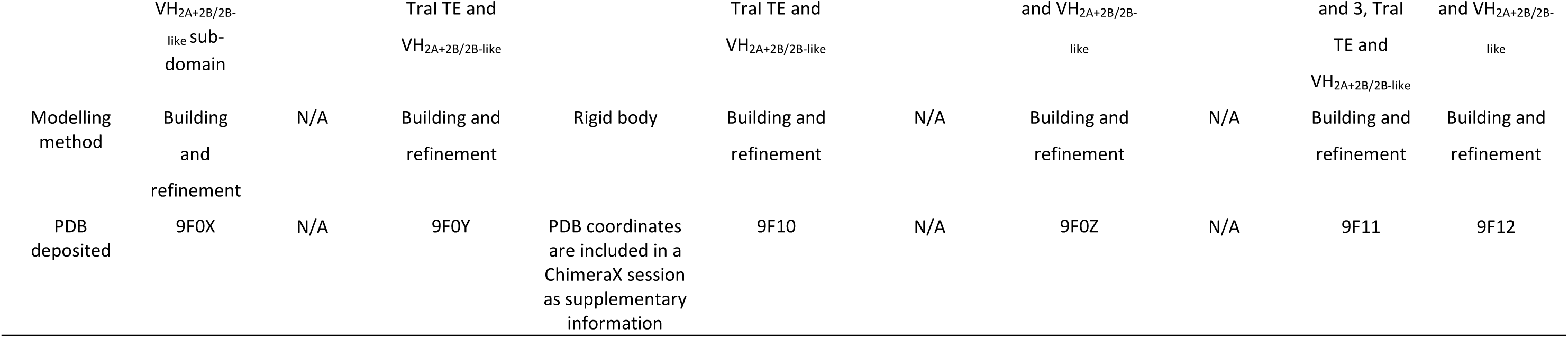
Cryo-EM data collection refinement and validations statistics and supplementary information. The supplementary information part of this table provides information on how each map was used, the part(s) that were built in each map when relevant, the method that was used to build the model from each map when relevant, and the PDB entry code of the corresponding model.

**ED Table 1b.**
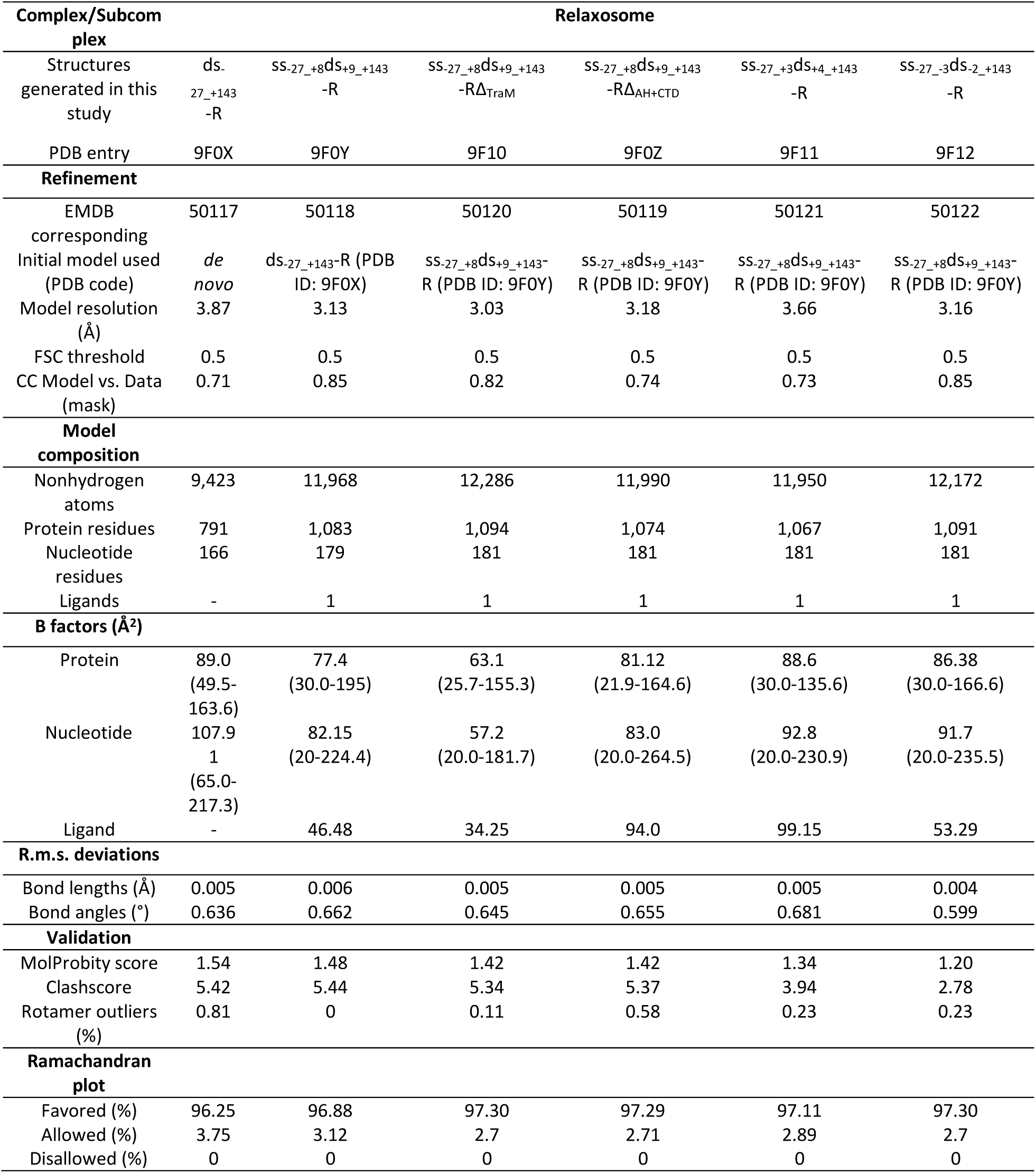
Model building and validation statistics.

**ED Table 2.**
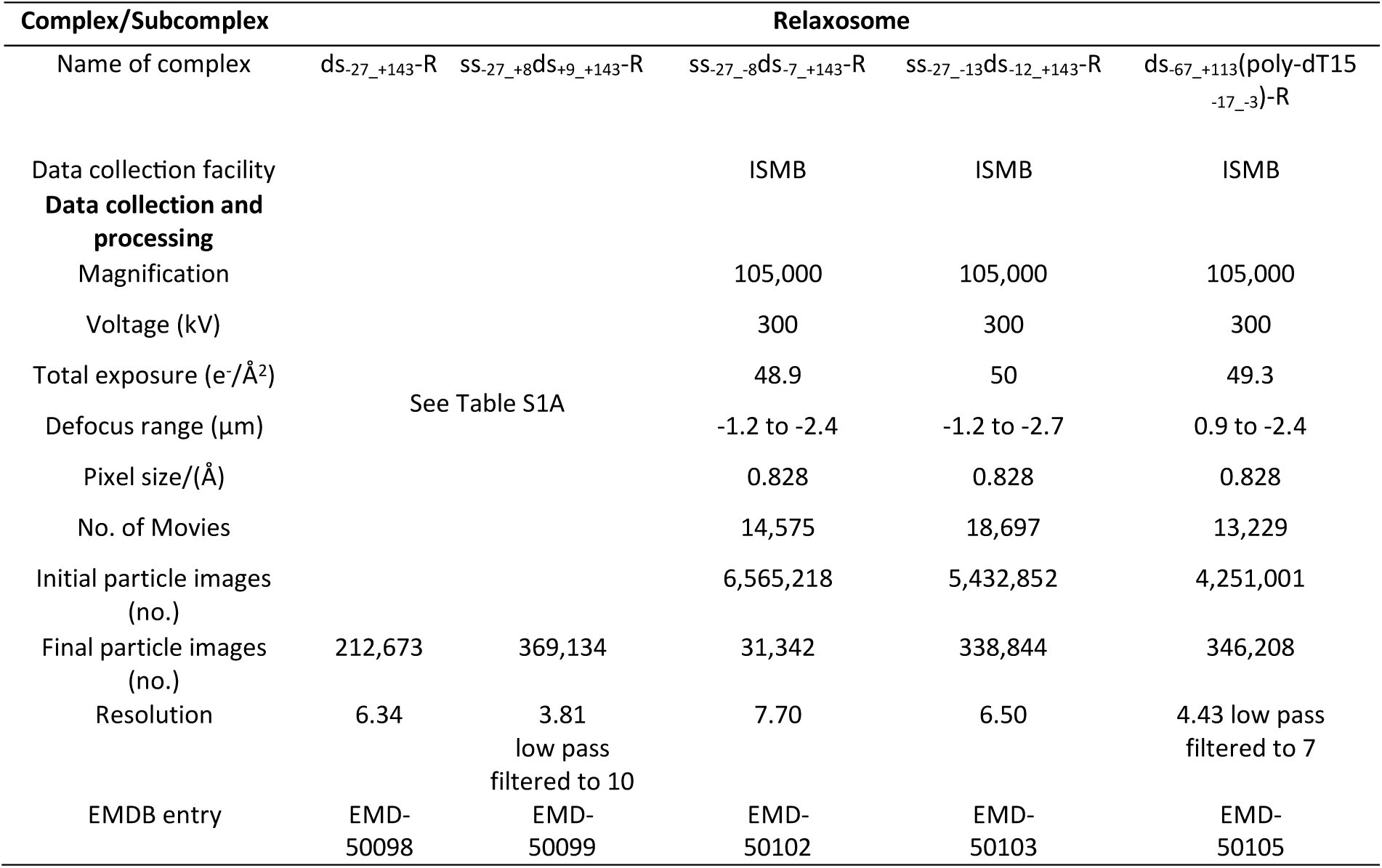
Cryo-EM data collection statistics for datasets used to generate 3D maps to illustrate DNA melting by the relaxosome.

| Complex/Subcomplex | Relaxosome |  |  |  |  |
| --- | --- | --- | --- | --- | --- |
| Name of complex | ds-27_+143-R | ss-27_+8ds+9_+143-R | ss-27_-8ds-7_+143-R | ss-27_-13ds-12_+143-R | ds-67_+113(poly-dT15-17_-3)-R |
| Data collection facility |  |  | ISMB | ISMB | ISMB |
| **Data collection and processing** |  |  |  |  |  |
| Magnification |  |  | 105,000 | 105,000 | 105,000 |
| Voltage (kV) |  |  | 300 | 300 | 300 |
| Total exposure (e<sup>-</sup>/Å<sup>2</sup>) |  |  | 48.9 | 50 | 49.3 |
| Defocus range (μm) |  | See Table S1A | -1.2 to -2.4 | -1.2 to -2.7 | 0.9 to -2.4 |
| Pixel size/(Å) |  |  | 0.828 | 0.828 | 0.828 |
| No. of Movies |  |  | 14,575 | 18,697 | 13,229 |
| Initial particle images (no.) |  |  | 6,565,218 | 5,432,852 | 4,251,001 |
| Final particle images (no.) | 212,673 | 369,134 | 31,342 | 338,844 | 346,208 |
| Resolution | 6.34 | 3.81<br>low pass<br>filtered to 10 | 7.70 | 6.50 | 4.43 low pass<br>filtered to 7 |
| EMDB entry | EMD-50098 | EMD-50099 | EMD-50102 | EMD-50103 | EMD-50105 |

**ED Table 3.** Functional analysis of *traI* mutants. Donor cells carrying the plasmids in the first column were mated with *E. coli* MS614 for 60 minutes. Average number of transconjugants obtained relative to donor cells is shown.

|  |  | <sup>a</sup>Conjugation frequency |  |  |
| --- | --- | --- | --- | --- |
|  | Variant of F *Tral* | Mean | SD | *P-value |
| **Donor *E. coli* MS411 [pOX38]** |  |  |  |  |
| pTrc99A | vector | 2.93 x 10<sup>-1</sup> | 0.1520 | n.a. |
| **Donor *E. coli* MS411 [pOX38Δ*tral*]** |  |  |  |  |
| <sup>b</sup>p99I<sup>+</sup> | wild-type | 1.11 x 10<sup>-3</sup> | 0.0007 | --- |
| **Donor *E. coli* MS411 [pOX38Δ*tral*]** |  |  |  |  |
| **DNA binding** |  |  |  |  |
| p99I<sup>+</sup>_Mut3 | R576A R605A | 5.89 x 10<sup>-4</sup> | 0.0003 | >0.9999 |
| p99I<sup>+</sup>_Mut4 | R576E R605E | 2.97 x 10<sup>-3</sup> | 0.0014 | >0.9999 |
| p99I<sup>+</sup>_Mut5 | S641E S696E | 8.45 x 10<sup>-1</sup> | 0.3002 | <0.0001 |
| p99I<sup>+</sup>_Mut6 | R694E S696E | 6.53 x 10<sup>-1</sup> | 0.1659 | 0.0001 |
| p99I<sup>+</sup>_Mut7 | S641E R694E S696E | 6.13 x 10<sup>-1</sup> | 0.1834 | 0.0002 |
| p99I<sup>+</sup>_Mut11 | R651E N799E | 8.20 x 10<sup>-4</sup> | 0.0005 | >0.9999 |
| **Tral-TraY** |  |  |  |  |
| p99I<sup>+</sup>_Mut12 | L646E R647E | 2.13 x 10<sup>-3</sup> | 0.0008 | 0.8778 |
| p99I<sup>+</sup>_Mut13 | L646R D648R | 3.74 x 10<sup>-2</sup> | 0.0158 | 0.0016 |
| **Tral-IHF** |  |  |  |  |
| p99I<sup>+</sup>_Mut14 | S678K S680R | 1.56 x 10<sup>0</sup> | 0.3140 | <0.0001 |
| p99I<sup>+</sup>_Mut15 | R673E T675E | 5.23 x 10<sup>-2</sup> | 0.0217 | 0.6636 |
|  | R673E T675E |  |  | 0.2139 |
| p99I<sup>+</sup>_Mut16 | Q677E | 2.01 x 10<sup>-1</sup> | 0.0106 |  |
| **TE-VH** |  |  |  |  |
| p99I<sup>+</sup>_Mut17 | S73E W659E | 1.24 x 10<sup>0</sup> | 0.6521 | 0.0181 |
| p99I<sup>+</sup>_Mut18 | S700E S701E | 9.51 x 10<sup>-1</sup> | 0.5485 | 0.0293 |

<sup>a</sup>Transconjugants/donor n=3; <sup>b</sup>n=4, \*adjusted P-values (ordinary one way anova, Holm-Šidák's multiple comparisons test comparing wild-type and mutants frequencies within respective groups), P-values <0.05 were considered significant

**ED Table 4.**
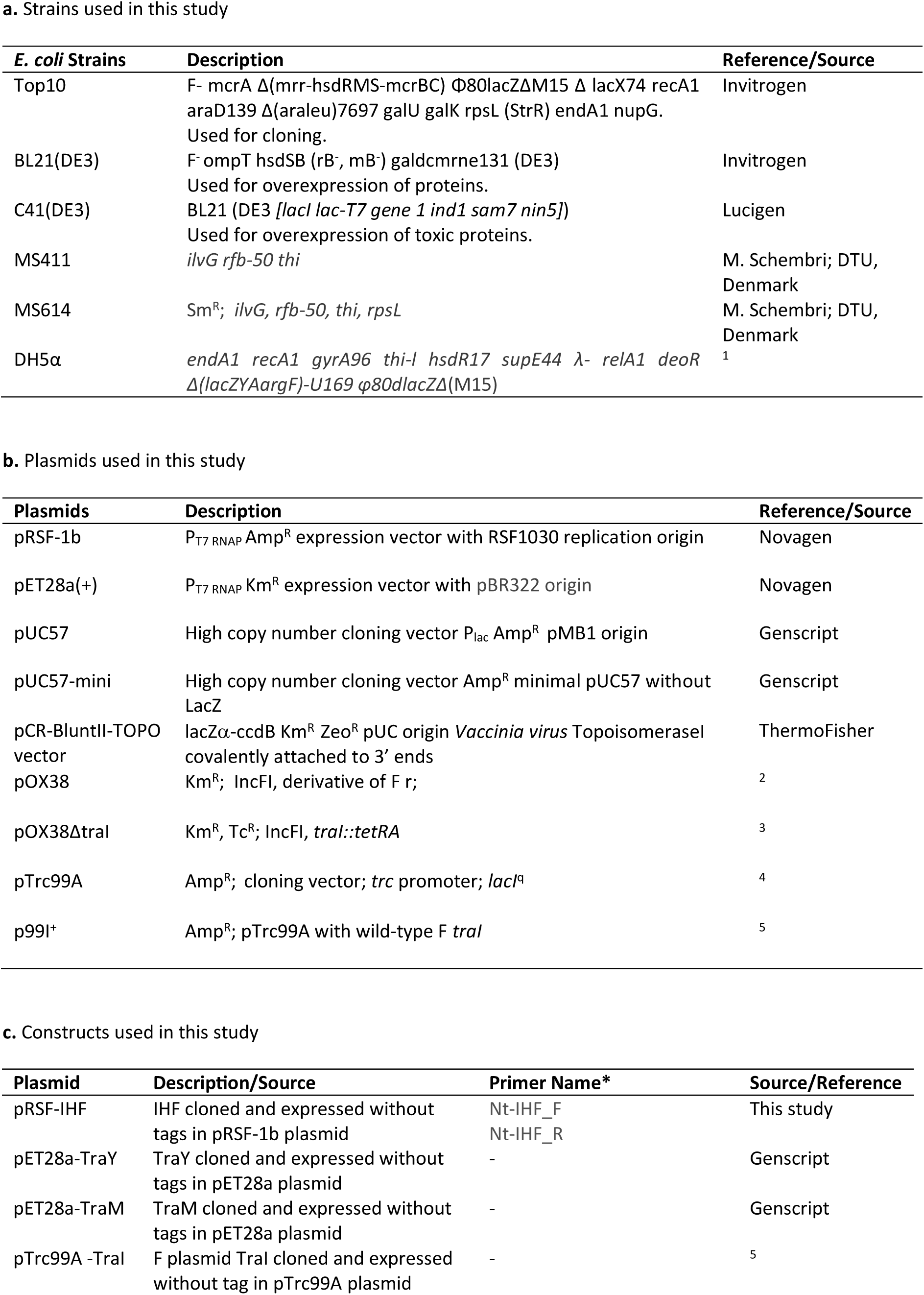
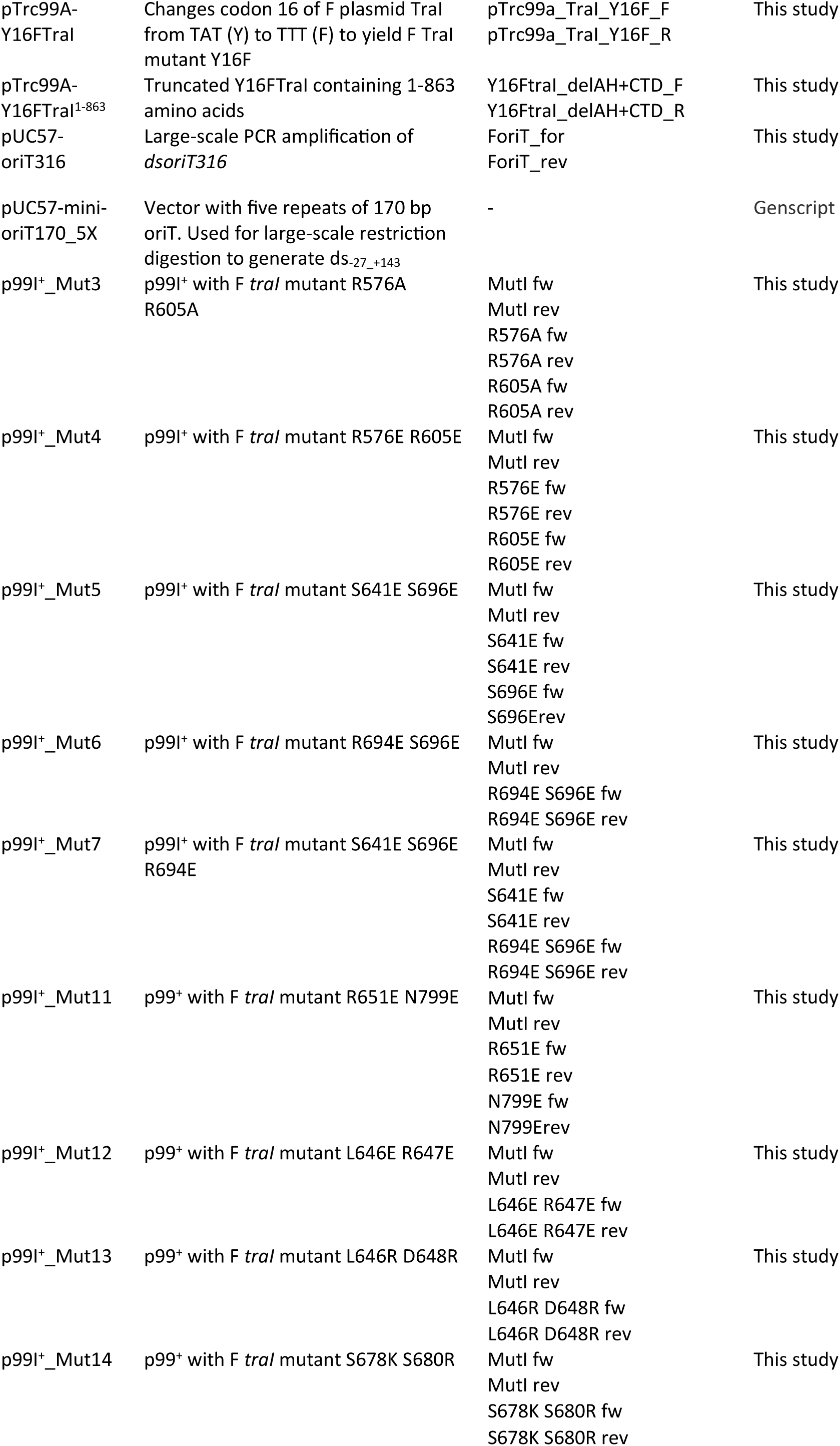
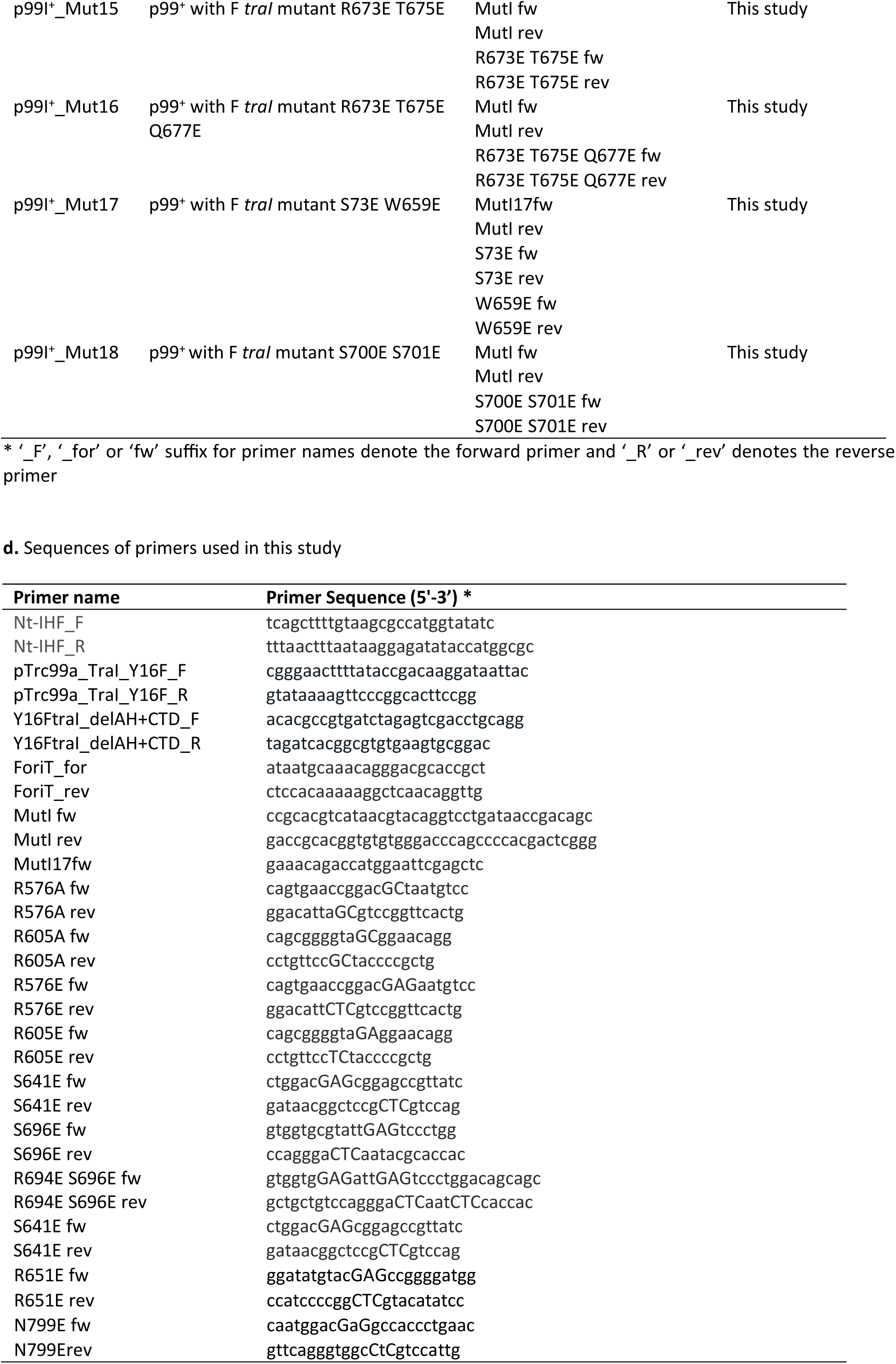
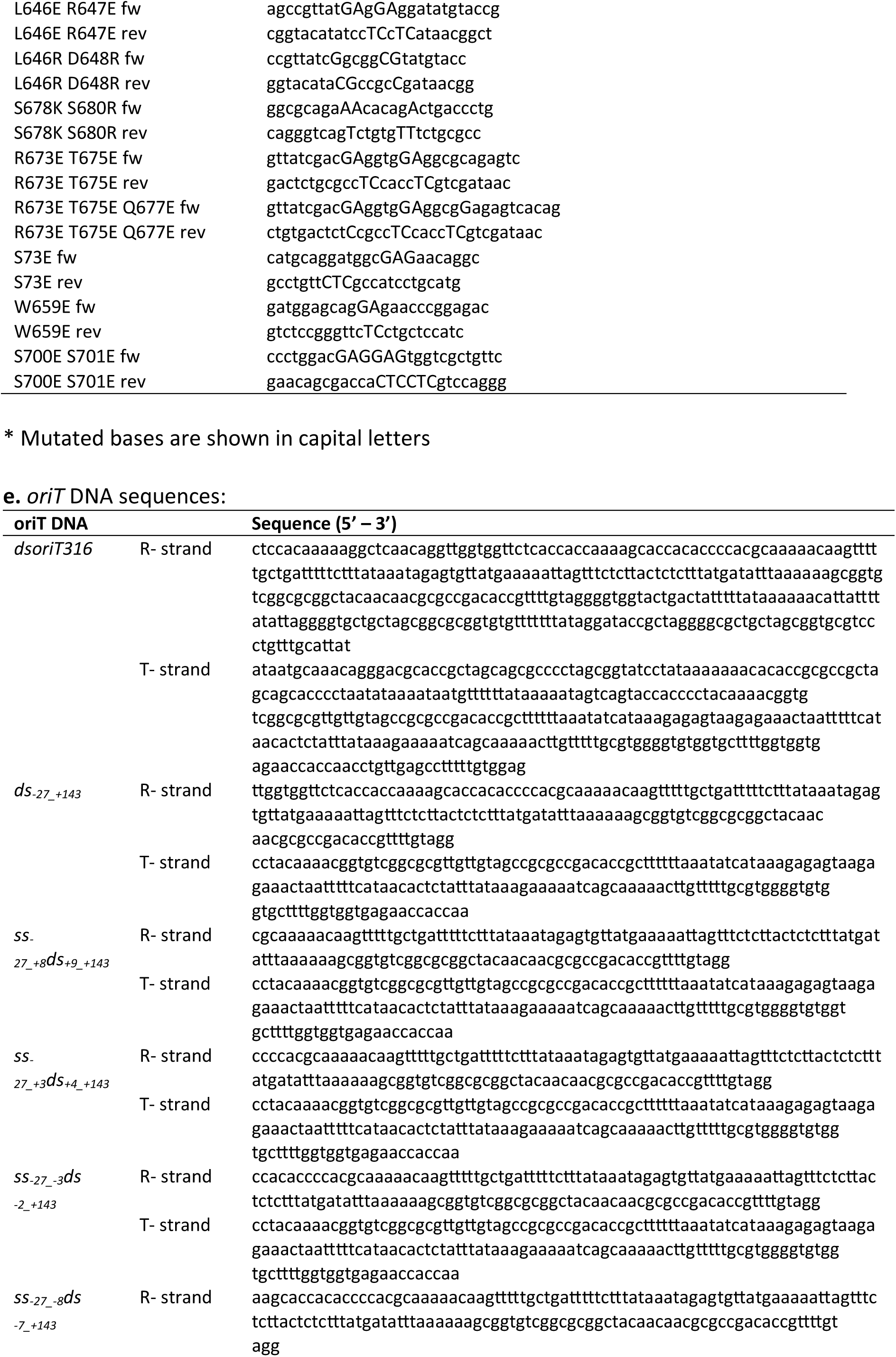
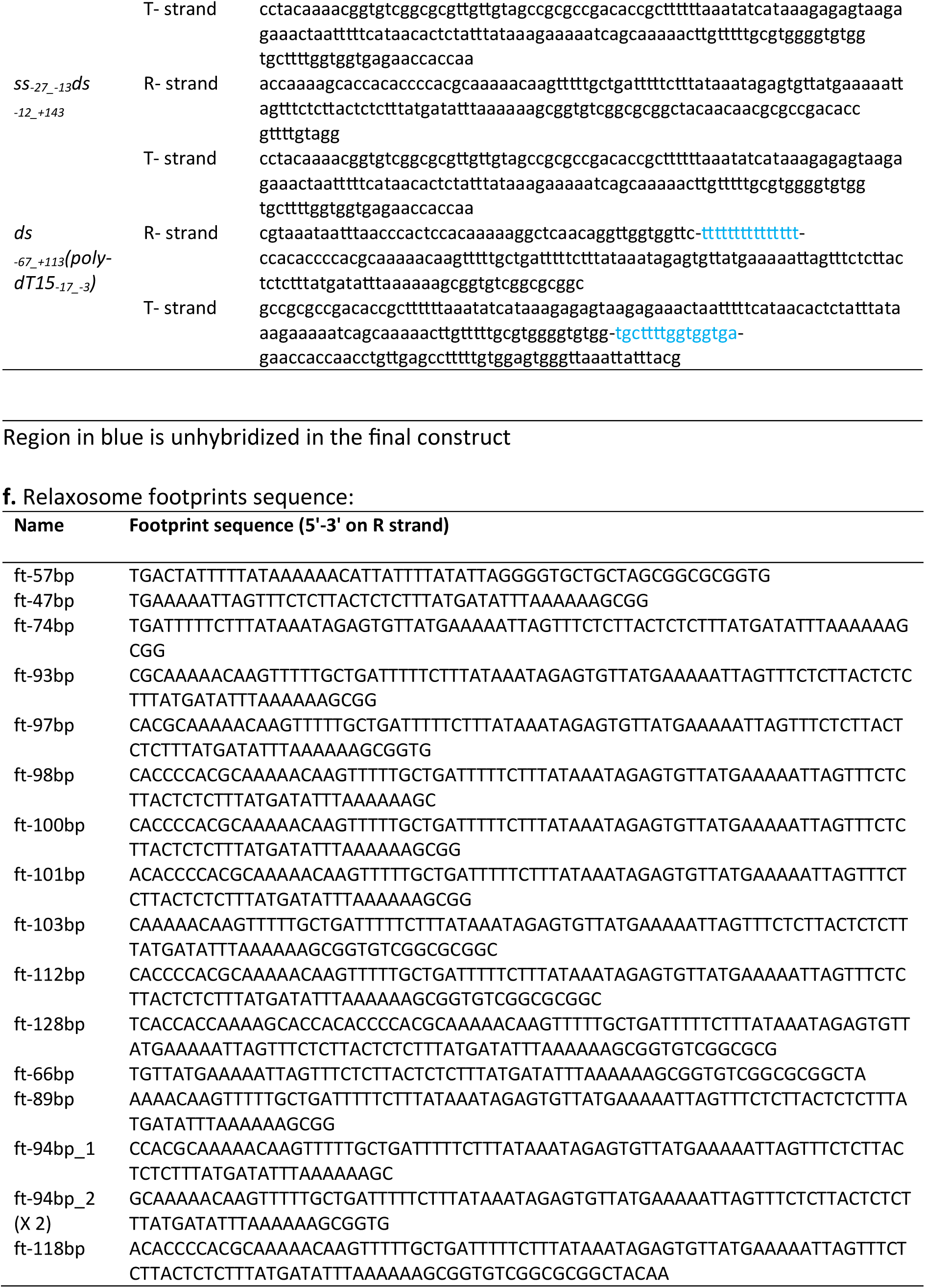
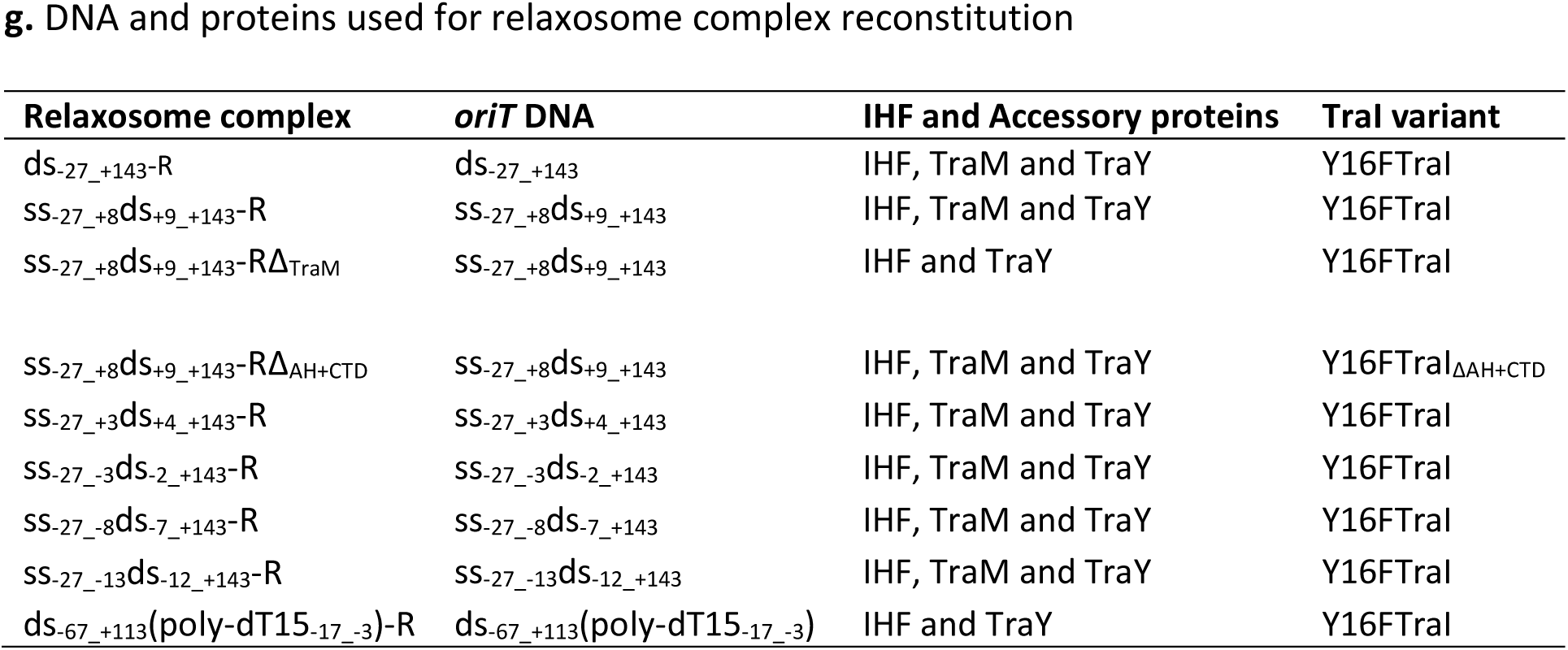
Strains, plasmids, constructs and oligonucleotides used in this study.

